# Joint Modeling of Effect Sizes for Two Correlated Traits: Characterizing Trait Properties to Enhance Polygenic Risk Prediction

**DOI:** 10.1101/2025.04.28.650892

**Authors:** Chi Zhang, Geyu Zhou, Tianqi Chen, Hongyu Zhao

## Abstract

Recent years have witnessed a surge in the development of innovative polygenic score (PGS) methods, driving their extensive application in disease prevention, monitoring, and treatment. However, the accuracy of genetic risk prediction remains moderate for most traits. Currently, most PGSs were built based on the summary statistics from the target trait, while many traits exhibit varied degrees of shared genetic architecture or pleiotropy. Appropriate leveraging of pleiotropy from correlated traits can potentially improve the performance of PGS of the target trait. In this study, we present PleioSDPR, a novel method that jointly models the genetic effects of complex traits to characterize conditions under which considering pleiotropy enhances polygenic risk prediction. PleioSDPR models the joint distribution of effect sizes across traits, allowing SNPs to be null for both traits, causal for only one trait, or causal for both traits, while accommodating region-specific genetic correlations and unequal heritability between traits. Through extensive simulations and real trait applications, we demonstrate that PleioSDPR improves prediction performance compared with several univariant and multivariate PGS methods, especially when there is no validation dataset. For example, by incorporating information from schizophrenia or leg fat-free mass, PleioSDPR effectively improves the prediction accuracy of bipolar disease (14.2% accuracy gain) and hip circumstance (20.65% accuracy gain), respectively. Moreover, our findings demonstrate that traits exhibiting high genetic correlations and heritability, and low overlapping sample sizes contribute more to the improvement of prediction accuracy of the target trait. Overall, our study highlights the potential of PleioSDPR to enhance the accuracy of genetic risk prediction by leveraging pleiotropy and considering a broader spectrum of traits and diseases. These findings contribute to the understanding of polygenic risk prediction and underscore the importance of incorporating pleiotropic information for improved utilization in disease prevention and treatment strategies.

## Introduction

Polygenic score (PGS) is formulated by aggregating the estimated effects of numerous genetic markers across the human genome to assess an individual’s genetic predisposition to complex traits or disorders. Recently, the significance of PGS has markedly increased as it has the potential to enhance the efficiency of population screening, refine diagnosis, and optimize disease treatment [1]. Additionally, the rapid advancement in genome-wide association studies (GWASs) has been pivotal in broadening the applications of PGS across diverse scientific fields [2,3].

However, a notable limitation of current PGS models is their primary focus on a single trait. In reality, genetic correlations exist among various complex human traits or disorders [4-8]. This shared genetic basis, known as pleiotropy [9], inspires the generation of PGS by considering multiple traits simultaneously. Even though multiple multivariate PGS models have been developed, most of them were designed to improve the prediction of a single trait in different populations. For example, PRS-CSx [10] is the extension of PRS-cs [11] to use a continuous shrinkage prior to a couple of genetic effects across populations, but it does not consider the genetic correlation between two traits. SDPRX [12] applies a Bayesian nonparametric prior to model different complex genetic architectures between two traits. However, it does not consider the heterogeneity of genetic correlation in different regions across the genome. JointPRS [30] extends PRS-CSx [10] to improve prediction by leveraging genetic correlations across more than two populations. However, it does not account for local genetic correlations and was not designed for modeling the joint distribution of complex traits within a single population, where trait-specific causal SNPs must be considered and usually a small proportion of SNPs are jointly causal between two traits, even when the overall genetic correlation is high.

Although some methods like PleioPred [13] and mtPGS [14] have been proposed to improve PGS predictions by incorporating data from genetically correlated traits, they also have limitations. PleioPred leverages pleiotropy based on the assumption that the true effect sizes follow a four-component normal distribution. However, PleioPred is limited to simple assumptions of the genetic architectures between traits and does not account for local genetic correlations between them. mtPGS models the joint distribution of effect sizes across two traits by assuming a constant genetic correlation across the genome. However, it does not account for local genetic correlations and does not explicitly model SNPs with null effects for both traits or trait-specific null effects. Thus, it is necessary to develop a new method to better incorporate information from correlated traits by fully examining the relationship between complex traits through local genetic heritability and local genetic correlation.

In pursuit of this goal, we introduce a novel method called PleioSDPR that accounts for genetically correlated traits by integrating GWAS summary statistics and LD matrix with effect sizes under a hierarchical Bayesian model. PleioSDPR characterizes the joint distribution of the effect sizes of a SNP in two traits as null, trait-specific, or shared with correlation. It assumes that the genome can be divided into different regions, each with different local heritability. Additionally, it allows for different local genetic correlations for regions having causal effects shared between two traits. We compared the performance of PleioSDPR with existing methods through extensive simulations and real data analysis, considering three different levels of sample overlap. Our results show that PleioSDPR may substantially improve the prediction accuracy over existing methods for the target trait when the paired trait has a larger heritability, high genetic correlation, and a low level of sample overlap.

## Methods

### Overview of PleioSDPR

We consider observations of complex trait Y_1_ with N_1_ samples and complex trait Y_2_ with N_2_ samples and assume that there are N_s_ samples overlapped between two traits. Let X_1_ and X_2_ be the standardized genotypes of the individuals for these two traits, so we have the following models for these two traits:

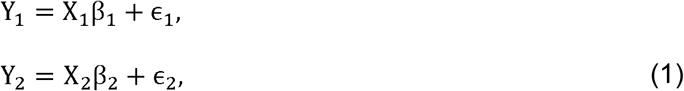

where β_1_ and β_2_ are the true effect size vectors for two traits, ϵ_1_ and ϵ_2_ are the vectors of residuals with 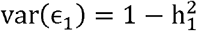 and 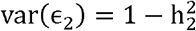, and 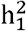 and 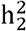 are the heritability of two traits, respectively. Since the covariance of residuals can be introduced by common environmental factors because of sample overlap [6,15,16], the residual covariance can also be called environmental covariance. For non-overlapped individuals, we simply assume that their environmental covariances are zero. However, if there are sample overlaps between two traits, we need to consider environmental covariance. We assume that the first N_s_ individuals overlapped for these two traits and let ρ_e_ be the environmental covariance, thus we could assume the residual covariance between these two traits as:

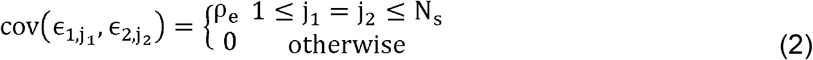

The likelihood model of PleioSDPR differs from previous multivariant models that aimed to integrate data from different populations. PleioSDPR focuses on connecting the marginal effect sizes in GWAS summary statistics for two traits within the same population to the true effect sizes, which needs to consider the environmental covariance within the same population due to sample overlapping. Thus, to accurately represent linkage disequilibrium (LD) and the environmental covariance due to sample overlapping, we construct the following multivariate normal distribution as the likelihood function, and the technical details are provided in the supplementary method.

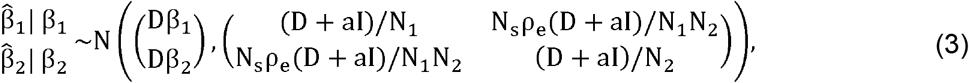

where 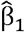 and 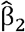 are the marginal effect sizes and D is the LD matrix. Same as SDPRX [12], this likelihood shrinks the off-diagonal covariance by a constant identity matrix to avoid the overestimation of effect sizes β_1_ and β_2_ for SNPs in high LD as a result of the mismatch between GWAS summary statistics and reference panel and we set *aN*_1_ and *aN*_2_ as 0.1. We set the overlap sample size N_s_ as input. If the accurate number of the overlap sample size can’t be obtained, we recommend using the minimal sample sizes of the individuals from the same cohort of two traits as the input of overlap sample size. Then combining with the slope and the intercept from the cross-trait LDSC [6], we can estimate 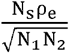 (Supplementary method). Except from the likelihood function, our prior also characterizes the joint distribution of the effect sizes for two traits in one population.

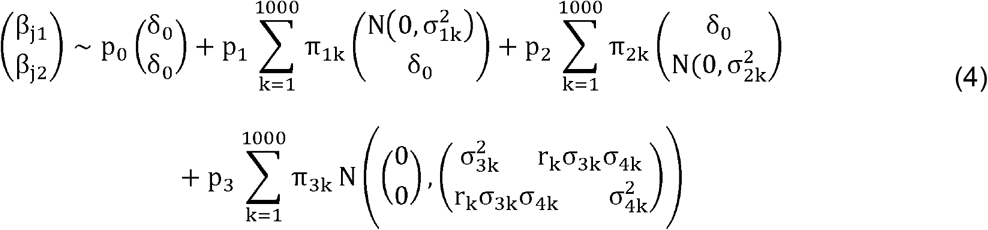

It assumes that a SNP for two traits can be classified into four possibilities: both null, trait specific, or shared with correlation, with the last three parts further divided into small components with different local heritability and local genetic correlations. In the previous analysis on exploring the genetic relationship between complex traits, we discovered that there were variations in both local heritability and local genetic correlations across regions and traits [5]. Thus, PleioSDPR allows the variances to be different for two traits and allows heterogeneity in genetic correlations between two traits in different chromosomal regions to better capture the information between complex traits.

Following SDPRX, PleioSDPR also uses a truncated stick-breaking process [12,17] to represent the probability of assignment for the second (trait1-specific), third (trait2-specific), and fourth terms (shared with correlation) and applies truncation by setting the maximum number of components of the mixture model as 1,000. Here we give the example of the fourth term (shared with correlation):

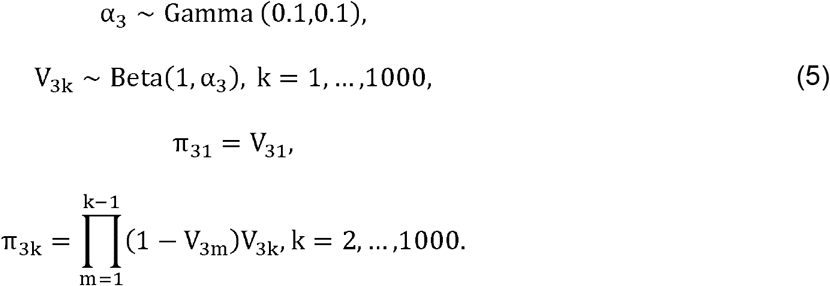

We apply Dirichlet distribution prior to the probability of each SNP belonging to each term:

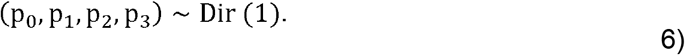

Priors of variance components for trait-specific terms are set to be hierarchical inverse gamma prior:

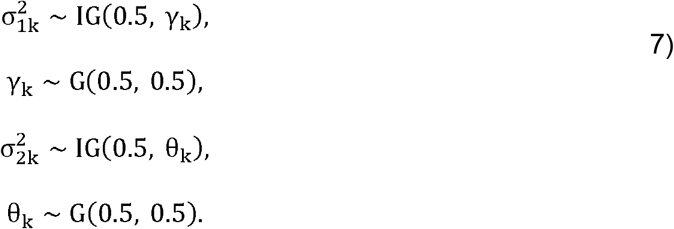

Besides, as suggested by Zhai et al. [18], we assume the prior of variance-covariance for the fourth term as hierarchical half-t prior:

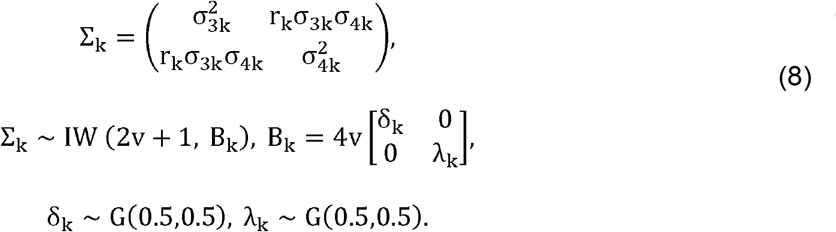

We partitioned the LD matrix into approximately independent LD blocks to reduce the computational burden [12,19] and adopted a Markov chain Monte Carlo (MCMC) algorithm based on the Gibbs sampler (supplementary method) to estimate the posterior effect sizes with 1,000 MCMC iterations and the first 200 iterations as the burn-in.

### Compared Methods

In our analysis, we evaluated the performance of PleioSDPR with four other approaches: LDpred2, PRS-CS, PRS-CSx, and SDPRX [10-12,20]. These methods fall into two primary categories. The first category includes univariate PGS models (LDpred2, PRS-CS) that use GWAS data from a trait as input. The second category includes multivariate PGS methods (PRS-CSx, SDPRx); compared to our model, PRS-CSx can be used as a reference when genetic correlations are not considered, while SDPRx accounts for genetic correlations but assumes they are constant across the genome.

When comparing the performance of different methods, we considered two distinct scenarios: one incorporating a validation dataset and another not. Without a validation set, the strategy of the linear combination of PGSs from multiple traits and tuning the methods’ parameters is infeasible. Thus, when there was no validation dataset, for LDpred2, PRS-CS, and PRS-CSx, we applied their auto versions (LDpred2-single, PRS-CS-single, PRS-CSx-single). SDPRX and PleioSDPR do not need to tune the parameters (SDPRx-single, PleioSDPR-single).

When there was a validation dataset, for all the methods a linear combination of PGSs from different traits was performed to derive the final score for the target trait (Ldpred2-mult, PRScs-mult, PRScsx-mult, SDPRx-mult, PleioSDPR-mult). For PRS-CSx and PRS-CS, the global shrinkage parameter was specified as {1e−06, 1e−04, 1e−02, 1, auto}. For LDpred2, we ran LDpred2-inf, LDpred2-auto, and LDpred2-grid and reported the best performance of the three options. The grid of hyperparameters of LDpred2-grid was set as non-sparse, *p* in a sequence of 17 values from 1e-05 to 1 on a log scale, and *h*^2^ within {0.7, 1, 1.4} of 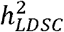.

### Simulations

In our simulation study, we utilized individual-level genotype data from unrelated, white British participants from the UK Biobank (UKB) [21]. We selected SNPs that were common between the UK Biobank, 1000 Genomes Phase 3, and HapMap3 datasets, resulting in a total of M=30,000 SNPs by selecting the first 3,000 SNPs from chromosomes 1 to 10. Our analysis included three sample sets. We randomly chose 20,000 individuals for set 1 and set 2 respectively that did not overlap with each other. Then we randomly selected 10,000 individuals from set 1 and set 2 respectively to construct set 3 to explore partial sample overlap scenarios. We assumed the underlying genetic architectures of the effect sizes for two traits included five parts, where 88% of the SNPs had zero effects on both traits, 2% of the SNPs had non-zero effects on only one trait, 1% of the SNPs had non-zero effects on both traits with high genetic correlation (r_1_ 0.9) and the remaining SNPs also had non-zero effects on both traits but with genetic correlation ranging from moderate to high (r_2_ 0.5,0.7,and 0.9). By incorporating SNPs with different levels of correlation across specific subsets of the genome, our simulation accounted for local genetic correlations rather than assuming a constant genetic correlation across the genome.

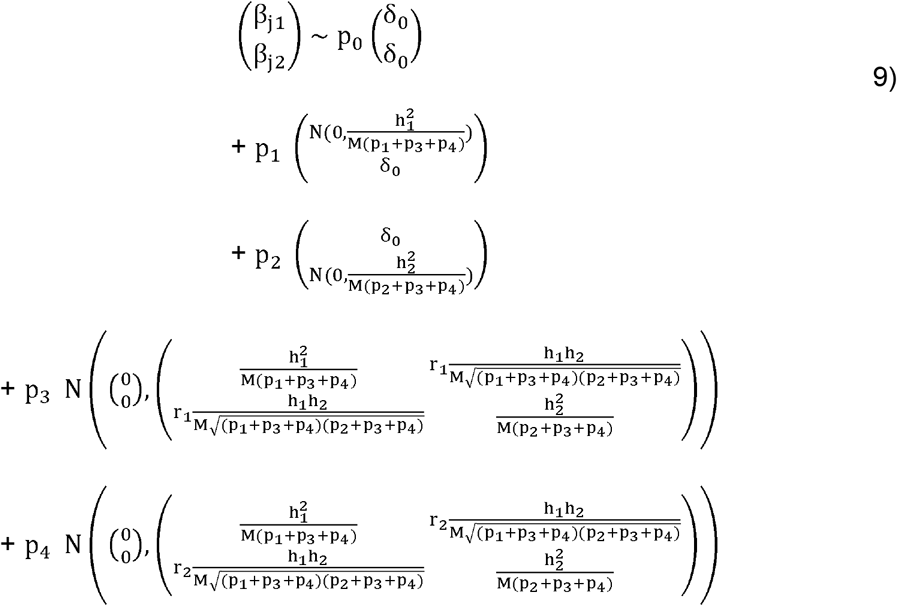

The heritability of the first trait was set to be h_1_^2^ =0.15 and the heritability of the second trait was set to be h_2_^2^ =0.3. When there were sample overlaps, we set the environmental covariance between two traits as 0.2. Then we applied GCTA [22] to generate continuous phenotypes and used PLINK 2.0 [23,24] to generate GWAS data using the training samples (set 1, set 2, or set 3). We replicated each simulation setting 10 times and evaluated the performance of different PGS methods using Pearson’s correlation.

We evaluated different PGS methods on three simulation scenarios with different levels of sample overlap. In the first scenario, set 1 and set 2 were used as independent training samples for two traits, ensuring no sample overlap. In the second scenario, set 1 and set 3 are chosen as the training samples, representing a scenario with partial sample overlap. Finally, in the last scenario, we selected set 1 to conduct GWASs for both traits, reflecting a scenario with complete sample overlap.

### Real data analysis

In real data analysis, we also tried to evaluate the performances of PleioSDPR under different levels of sample overlap (No sample overlap, partial sample overlap, and complete sample overlap). We obtained GWASs from various consortia including the Genetic Investigation of ANthropometric Traits (GIANT), DIAbetes Genetics Replication And Meta-analysis (DIAGRAM), and the Psychiatric Genomics (PGC) Consortium. To account for different levels of sample overlap comprehensively, we conducted GWASs utilizing individual-level data from the UKB dataset. We chose 381,924 unrelated White British individuals and selected SNPs overlapped with the Hapmap3 dataset. We conducted BOLT-LMM [25] to generate GWASs, adjusting for covariates including sex, age, age squared, sex by age interaction, sex by age squared interaction, genotyping array, and the first 20 genotype principal components.

In the scenario where there was no sample overlap between two traits, we compared PGS methods across three trait pairs — waist circumference and weight, hip circumference and leg fat-free mass, and type 2 diabetes (T2D) and waist-hip ratio (WHR) — which not only exhibit high global genetic correlations but also contain multiple regions with significant local genetic correlations of varying magnitudes. The GWASs of waist circumference, hip circumference, T2D, and WHR were obtained from publicly available datasets. We generated the GWASs of weight and leg fat-free mass using about 80% selected unrelated White British individuals in UKB. We evaluated the performance of these methods within the unrelated European population of the UK Biobank, specifically individuals born in England (identified by Field 1647 coded as 1), who were not included in the training dataset used to generate GWASs. Other unrelated European individuals in UKB that were not included in both training and testing datasets were used for parameters tuning and linear combination, which were divided into three validation datasets with different sample sizes. We validated each PGS method in different validation datasets and assessed their performance on the same testing dataset obtaining three *R*^2^ values. The mean of these *R*^2^ values served as our final evaluation metric. Sample sizes for the testing and validation datasets are summarized in Supplementary Table 1.

A summary of the sample sizes, heritability, genetic correlation and the number of regions with significant local genetic correlation for these traits is presented in Table 1. Heritability and genetic correlation were calculated using LDSC [6,26]. We chose these three trait pairs because they have unbalanced heritability and high genetic correlation.

**Table 1:**
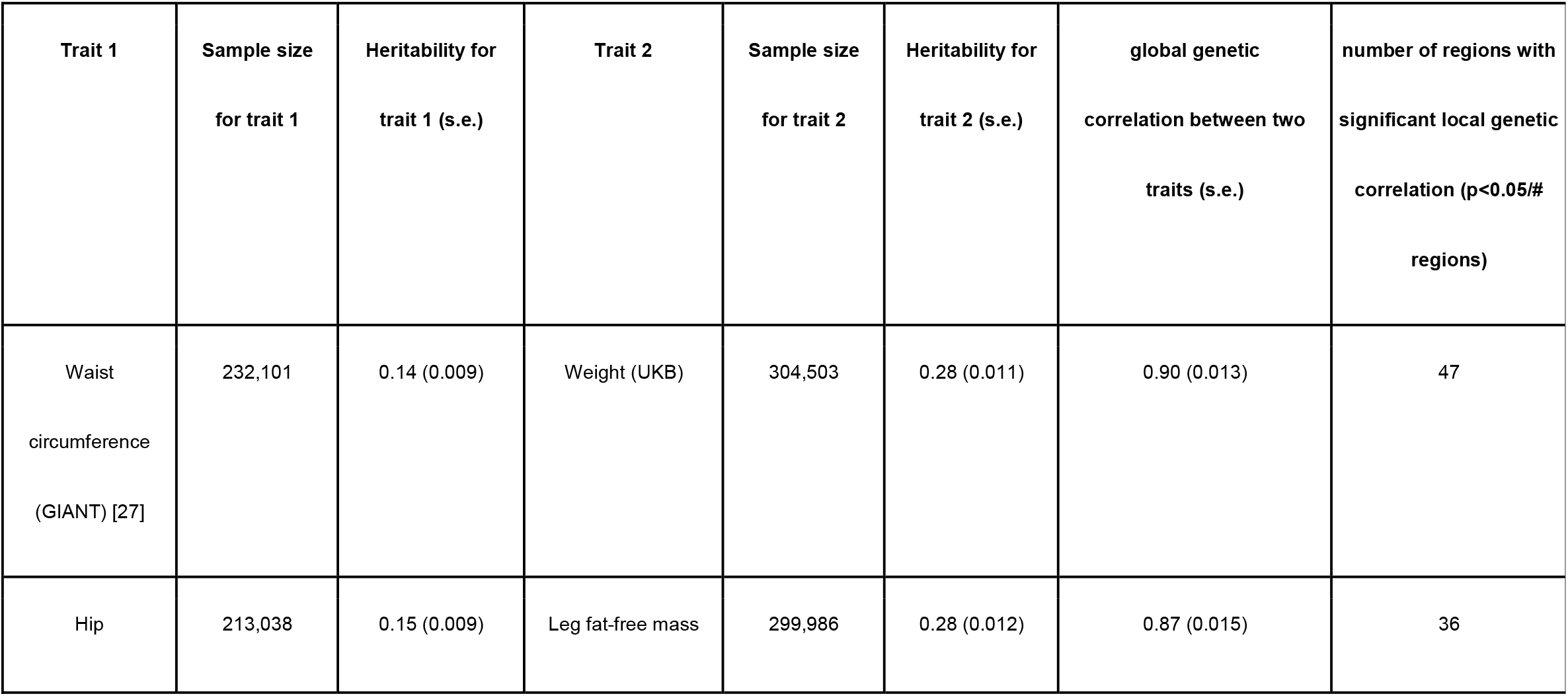
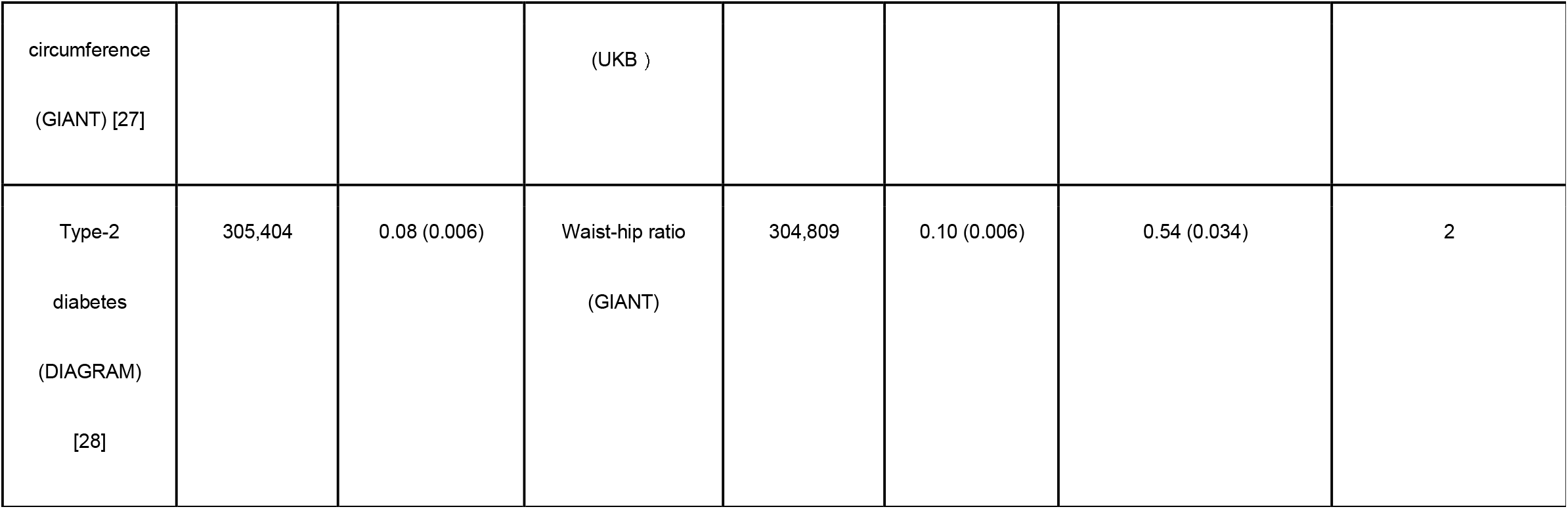
Trait-pairs for real data analysis without sample overlap. We provide the sample sizes, heritability, and source of GWASs for these six traits. We also show the genetic correlation between each trait pair. Heritability and genetic correlation were calculated from LDSC [8, 26]. The number of regions with significant local genetic correlation were calculated from SUPERGNOVA [4].

In the scenario with partial sample overlap between two traits, we continued our comparison for waist circumference and weight, as well as hip circumference and leg fat-free mass. The GWASs of these four traits were conducted using individual-level data from the UKB. Specifically, we utilized the same GWASs for weight and leg fat-free mass as those from the real data analysis with no sample overlap. For waist circumference and hip circumference, we generated GWASs using approximately half of the individuals used for the weight and leg fat-free mass GWASs. Additionally, we introduced a new trait pair, schizophrenia (SCZ) and bipolar disorder (BIP), for which GWASs were obtained from the PGC consortium. Same as waist circumference and weight, and hip circumference and leg fat-free mass, BIP and SCZ also have several regions with significant different local genetic correlations. The testing dataset and validation datasets remained consistent with the previous real data analysis. However, due to the limited number of cases for BIP and SCZ in the UKB, we only considered two validation datasets for this trait pair. Sample sizes for the testing and validation datasets are summarized in Supplementary Table 2. A summary of the sample sizes of each trait, overlapping sample size, heritability, global genetic correlation and the number of regions with significant local genetic correlation for these traits is provided in Table 2.

**Table 2:**
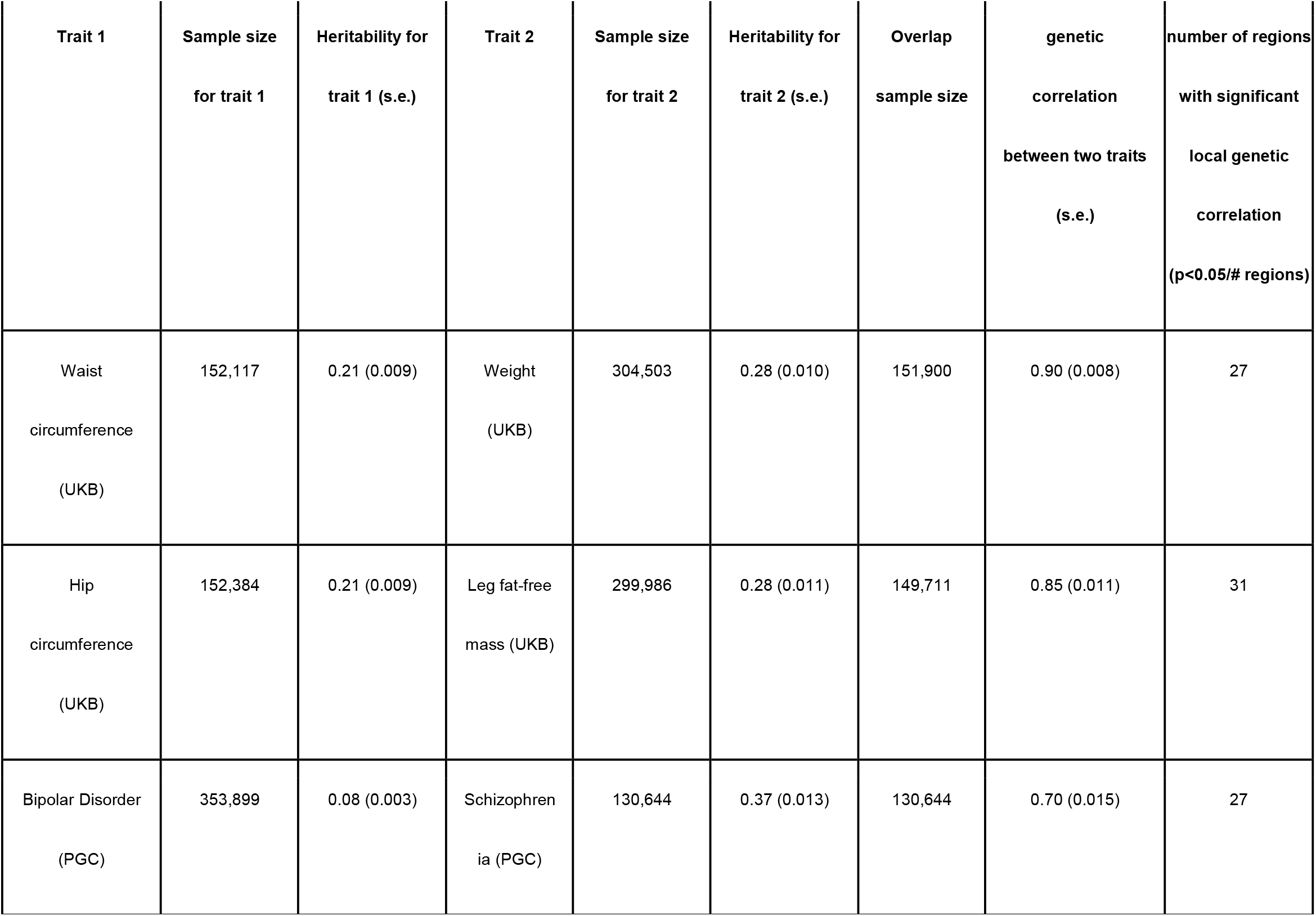
Trait-pairs for real data analysis with partial sample overlap. We provide the sample sizes of each trait, overlapped sample sizes, heritability, and source of GWASs for these six traits. We also show the genetic correlation between each trait pair. Heritability and genetic correlation were calculated from LDSC [8, 26]. The number of regions with significant local genetic correlation were calculated from SUPERGNOVA [4].

When the samples of two traits completely overlapped with each other, we only did our comparison for waist circumference and weight, as well as hip circumference and leg fat-free mass. The GWASs of these four traits were conducted using individual-level data from the UKB with the same sample sizes. We only considered the situation when there was no validation dataset. The testing dataset remained with the same as the previous real data analysis. A summary of the sample sizes of each trait, overlapping sample size, heritability, global genetic correlation and the number of regions with significant local genetic correlation for these traits is provided in Table 3.

**Table 3:** Trait-pairs for real data analysis with complete sample overlap. We provide the sample sizes of each trait, overlapped sample size, heritability, and source of GWASs for these six traits. We also show the genetic correlation between each trait pair. Heritability and genetic correlation were calculated from LDSC [8, 26]. The number of regions with significant local genetic correlation were calculated from SUPERGNOVA [4].

| Trait 1 | Sample size<br>for trait 1 | Heritability<br>for trait 1<br>(s.e.) | Trait 2 | Sample size<br>for trait 2 | Heritability<br>for trait 2<br>(s.e.) | Overlap<br>sample size | genetic correlation<br>between two traits<br>(s.e.) | number of regions with<br>significant local genetic<br>correlation ($p < 0.05/\#$<br>regions) |
| --- | --- | --- | --- | --- | --- | --- | --- | --- |
| Waist<br>circumference<br>(UKB) | 304,503 | 0.21 (0.008) | Weight<br>(UKB) | 304,503 | 0.28 (0.010) | 304,503 | 0.90 (0.008) | 31 |
| Hip circumference<br>(UKB) | 299,986 | 0.21 (0.008) | Leg fat-free mass<br>(UKB ) | 299,986 | 0.28 (0.011) | 299,986 | 0.85 (0.011) | 36 |

## Results

### Simulations

We began by evaluating the predictive performance of each method through simulations across multiple genetic correlations and levels of sample overlap. We considered five methods: PleioSDPR, SDPRX [12], PRS-CSx [10], PRS-CS [11], and LDpred2 [20]. LDpred2 and PRScs-CS are univariate methods that utilize summary statistics from a single trait, while PleioSDPR, SDPRX, and PRS-CSx are multivariate methods that jointly incorporate GWAS summary statistics from multiple traits. For simulation studies, we used genotype data from unrelated White British individuals from UKB. We selected a total of 30,000 SNPs, with 3,000 SNPs from each of chromosomes 1 to 10, respectively. The training cohort comprised 20,000 individuals for both traits. We considered three levels of sample overlap: no sample overlap, partial sample overlap (with an overlapped sample size of 10,000), and complete sample overlap. The validation and test datasets each consisted of 10,000 individuals. The genetic architecture simulated for the two traits was detailed in the Methods section, including unbalanced heritability (0.15 and 0.3) and different values of genetic correlations. Each simulation setting was repeated 10 times. We first generated summary statistics for two traits without sample overlap and used the 1KG phase 3 data [29] as the reference panel to estimate the LD matrix. We then evaluated the performance of each method using the square of the Pearson correlation of PGS and simulated phenotype in the independent testing dataset. In cases where a separate validation dataset was available, we used it to tune the parameters for LDpred2, PRS-CS, and PRS-CSx. We also performed linear combinations of the PGSs of the two traits. When no validation dataset was present, we applied the auto version of LDpred2, PRScs, and PRScsx, as well as the original versions of SDPRX and PleioSDPR, as parameter tuning was unnecessary for these methods.

Figure 1 presents the outcomes of simulations conducted for scenarios where two traits do not share samples. We first compared the performances of PGS methods when there was no validation dataset (with the “_single” suffix). PleioSDPR consistently outperformed the other PGS methods across all genetic correlations, particularly for traits with lower heritability. The results also reveal that the larger the genetic correlation between the two traits, the more accurate the prediction of PleioSDPR. We also observed that PleioSDPR, which modeled local genetic correlation, outperformed SDPRx which assumed a constant genetic correlation, and PRS-CSx which did not account for genetic correlation. This finding underscores the importance of modeling local genetic correlations, as accounting for region-specific genetic architectures enables more accurate identification of trait-specific and shared genetic effects, ultimately leading to improved polygenic score prediction. For methods with “_mul” suffix which means the inclusion of a validation dataset, there is a notable increase in accuracy for all methods except PleioSDPR. This suggests that these methods benefit significantly from the validation dataset. In contrast, PleioSDPR maintains a relatively stable accuracy no matter whether the validation dataset exists or not and consistently performs the best among all the compared methods. This indicates that PleioSDPR can learn sufficient information directly from the GWAS data, making the validation dataset less necessary for its performance improvement compared to the other methods. When analyzing the simulation results under conditions of partial sample overlap (Supplementary Figure 1), we found that PleioSDPR maintains its superiority over alternative methods in scenarios without a validation dataset. However, this advantage is less obvious in cases with no sample overlap between two traits (Supplementary Figure 1). With the introduction of a validation dataset, different methods yielded similar performance, with PleioSDPR consistently achieving or matching the highest levels of predictive accuracy.

**Figure 1:**
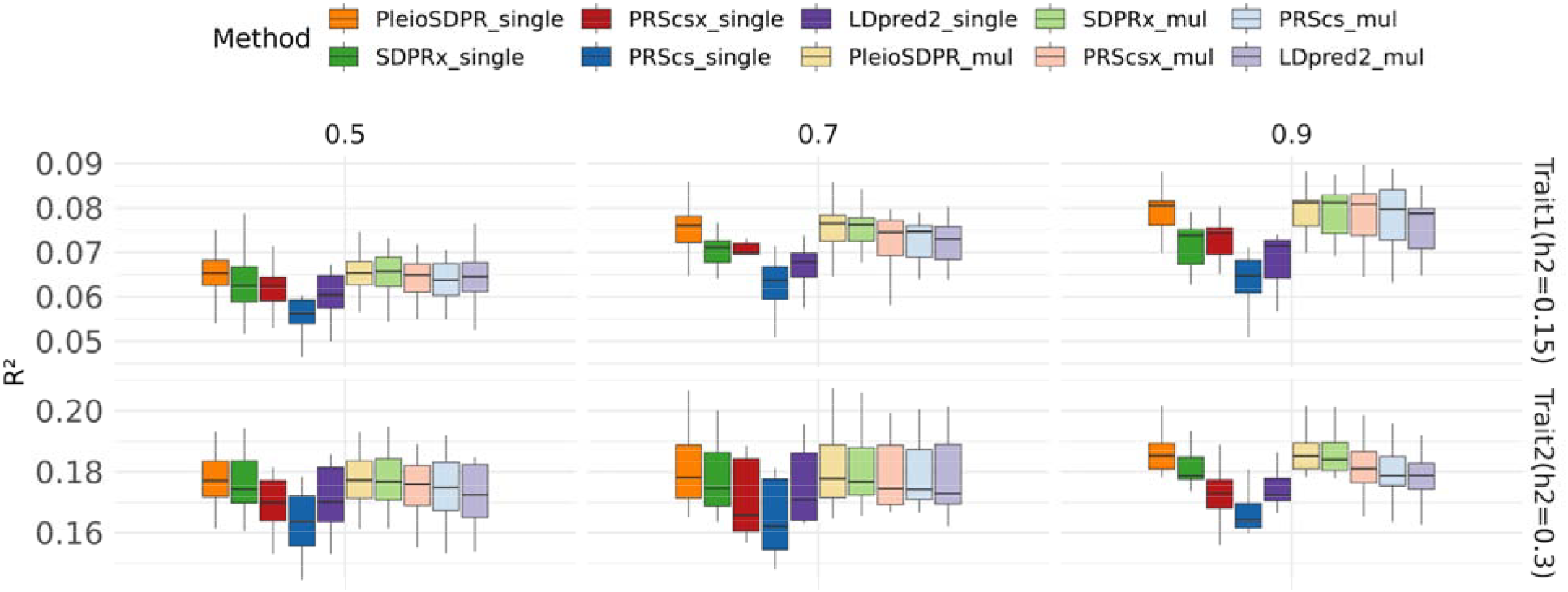
Simulation results when there is no sample overlap. We compare the methods considering with validation dataset and without validation dataset. When there is no validation dataset, we name them as PleioSDPR_single, SDPRx_single, PRScsx_single, PRScs_single and LDpred2_single. When there is a validation dataset, we name them PleioSDPR_mul, SDPRx_mul, PRScsx_mul, PRScs_mul, and LDpred2_mul.

When the samples of two traits completely overlapped PleioSDPR performed exclusively the best when the genetic correlation of two traits was high (0.9) and the prediction was on the first trait with lower heritability in the absence of the validation dataset (Supplementary Figure 2). Given that the two traits fully overlap with each other, information obtained from correlated traits due to increased sample size is not possible; rather, the method benefits from information from the traits with higher heritability. This suggests that to demonstrate optimal performance, there must be enough information from the paired correlated traits that can be borrowed to help improve the prediction of the target trait. In simulations of complete sample overlap, PleioSDPR does not outperform all competing methods; however, it at least shows performance comparable with the other methods.

### Prediction performance in real data analysis

Subsequently, we evaluated PleioSDPR’s performance through real data analysis across various trait pairs. We examined the effect of varying degrees of sample overlap and evaluated the performance of these methods both with and without validation data. We used validation datasets of different sample sizes to tune parameters and evaluated the performance on the same testing dataset. We used the mean of R-squared as the measurement of prediction accuracy.

We first selected three pairs of traits with no overlapping samples: Waist Circumference and Weight, Hip Circumference and Leg Fat-Free Mass, and T2D and WHR. PleioSDPR demonstrates the best performance across all six traits (Figure 2). Within these trait pairs, waist circumference, hip circumference, and T2D exhibit lower heritability compared to their auxiliary traits, and PleioSDPR demonstrated more pronounced improvement for these lower-heritability traits. PleioSDPR outperformed SDPRx and PRS-CSx, both of which did not account for local genetic correlations. For all three trait pairs, local genetic correlation analyses revealed multiple regions with significant and varying local genetic correlation values. These findings emphasize that incorporating heterogeneous local genetic correlations into modeling can improve polygenic score prediction, particularly for traits with lower heritability. When a validation dataset is used for parameter tuning (Supplementary Figure 3), PleioSDPR still performs best or second-best with comparable performance to other PGS methods.

**Figure 2:**
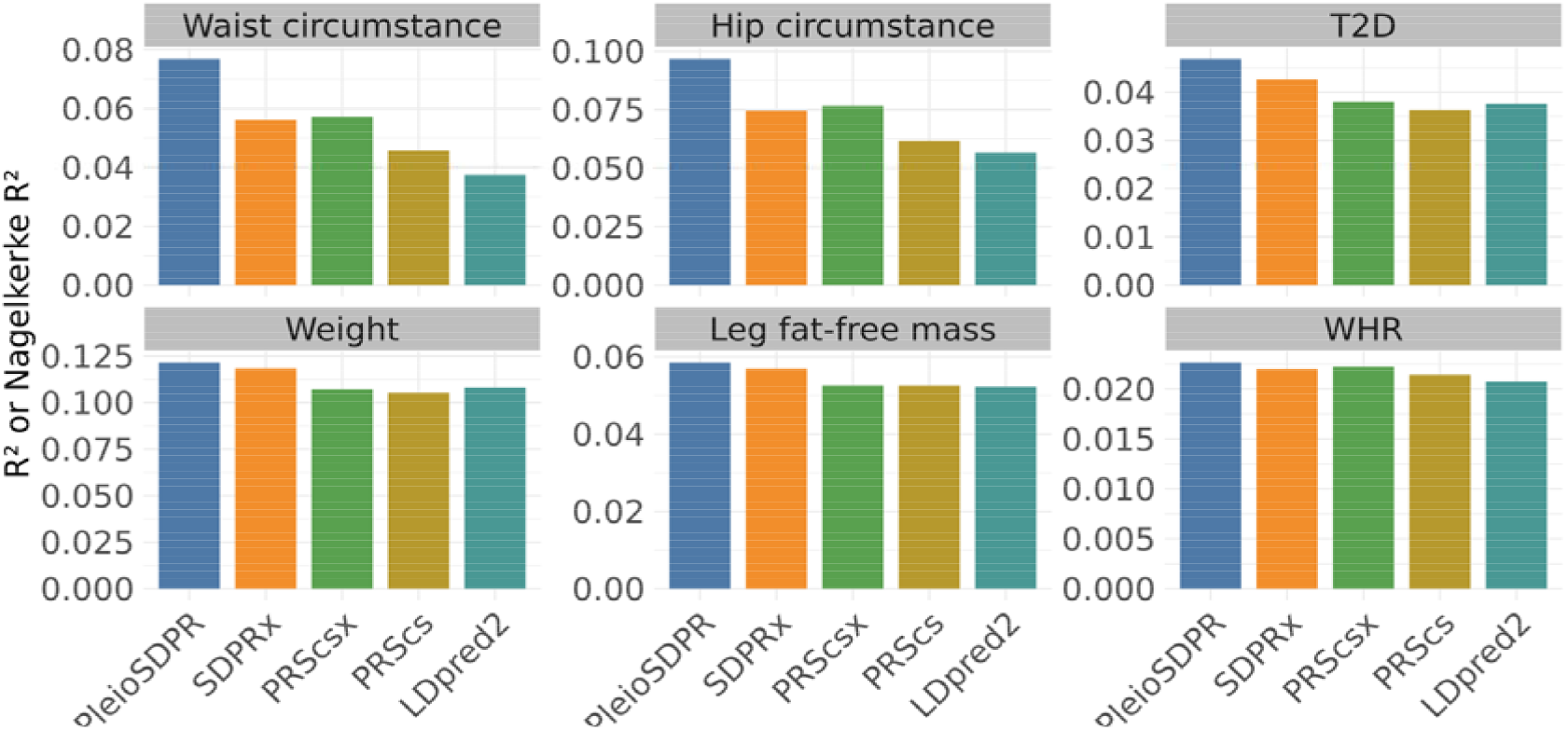
Prediction performance of different methods for three trait pairs that have no sample overlap when there is no validation dataset.

Environmental covariance is important when there is sample overlap between pairs of traits. To investigate the impact of sample overlap, we generated GWAS summary statistics in UKB for Waist Circumference and Weight, Hip Circumference, and Leg Fat-Free Mass. Additionally, BIP and SCZ were also considered under the condition of partial sample overlap. We first evaluated the performance of PleioSDPR with and without the consideration of environmental covariance. The comparison indicates that the inclusion of the estimation of environmental covariance enhances PleioSDPR’s performance (Supplementary Figures 4 and 6). We next compared the performance of PleioSDPR with other methods when there is sample overlap. In the absence of a validation dataset, PleioSDPR exhibits the best performance in predicting Waist Circumference, Hip Circumference, BIP, and SCZ while ranking second in accuracy for Weight and Leg Fat-Free Mass (Figure 3). This pattern mirrors the observations made in the simulation scenario without sample overlap, where PleioSDPR demonstrates a predictive advantage with traits that have lower heritability.

**Figure 3:**
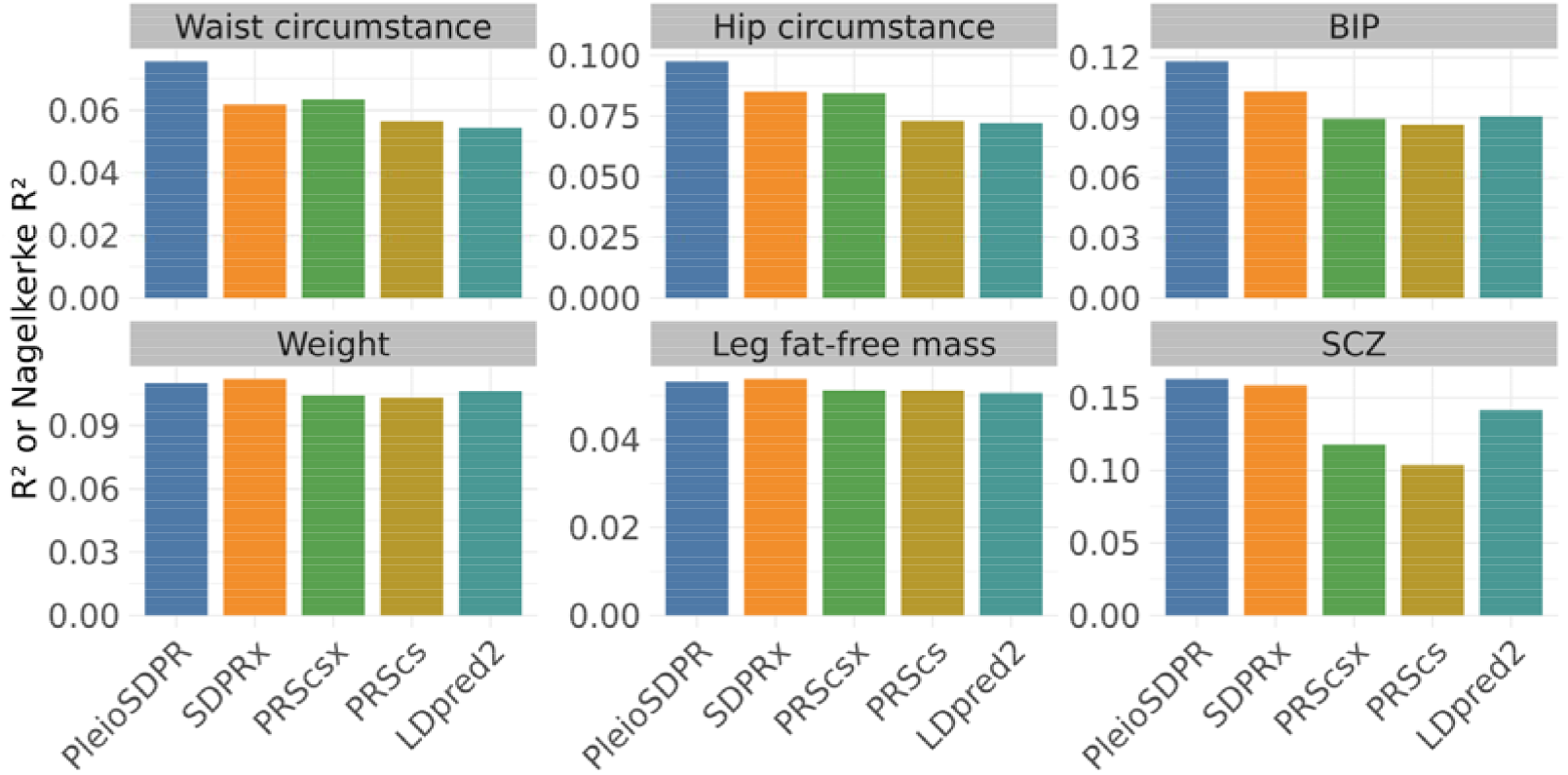
Prediction performance of different methods for three trait pairs that have partial sample overlap when there is no validation dataset.

When incorporating a validation dataset (Supplementary Figure 5), PleioSDPR remains the top-performing method for Waist Circumference, Hip Circumference, BIP, and SCZ, and ranks as the second most accurate for the remaining traits. However, across all traits, PleioSDPR’s performance is comparable with SDPRX when the validation dataset is considered. When the samples for two traits were entirely overlapping, we focused on two specific pairs: Waist Circumference and Weight, and Hip Circumference and Leg Fat-Free Mass. To ensure complete sample overlap among these trait pairs, we utilized the same cohort from the UK Biobank to generate GWASs for these traits. Our findings indicate that the performance of all methods is comparable across all traits, regardless of whether a validation dataset is considered. In this context, no single method distinctly outperforms the others (Supplementary Figures 7 and 8).

## Discussion

We have introduced a novel PGS model, PleioSDPR, designed to improve the prediction accuracy of the target trait by leveraging genetically correlated traits. PleioSDPR offers flexibility in modeling the genetic architecture between two complex traits, accounting for variations in local genetic heritability and local genetic correlations across different regions. Moreover, PleioSDPR considers the environmental covariance introduced by sample overlap by modifying the likelihood function that can be used in the MCMC algorithm. Notably, PleioSDPR eliminates the need for parameter tuning and maintains optimal performance when there is no validation dataset.

Building on the Bayesian nonparametric prior concept from SDPRX, PleioSDPR further explores the shared genetic architecture between the target trait and correlated traits by classifying SNPs into different categories according to their effect sizes. PleioSDPR assumes that SNPs may either be non-causal for both traits, causal for only one, or causal for both. When SNPs are categorized as causal for at least one trait, they are further divided into subcategories with different variances or covariances. Besides, by addressing the limitations of SDPRX, PleioSDPR allows for variation in the variance of SNPs that are shared between traits and accommodates different local genetic correlations within different subcategories. This method of SNP categorization enables PleioSDPR to apply shrinkage to the estimated effect sizes across different SNP categories, ensuring adequate shrinkage for small effects while preventing over-shrinkage of large effects, particularly in SNPs that are causal for both traits.

In this study, we evaluated the performance of PleioSDPR through both simulations and real data analyses. We found that selecting GWAS data from correlated traits that not only share similarities with the target trait but also provide complementary genetic information is critical for improving prediction accuracy. Specifically, appropriately leveraging pleiotropy—by utilizing shared genetic effects, regions of overlapping causality, and supplementary heritability information—can substantially enhance the performance of polygenic risk scores, particularly when the auxiliary trait exhibits high genetic correlation, substantial heritability, and minimal sample overlap with the target trait. Our simulation settings and real data applications considered scenarios involving two correlated traits with unbalanced heritability, moderate to high genetic correlation, varying degrees of sample overlap and multiple regions with different significant local genetic correlations. These results highlight that incorporating pleiotropic information can meaningfully improve genetic risk prediction for complex traits. In particular, modeling heterogeneous genetic correlations across genomic regions better captures the underlying genetic architecture, where local correlation patterns vary across the genome. However, introducing region-specific parameters also increases estimation variance. Without sufficient information borrowing across traits and regions, the additional variance from modeling multiple local correlations may offset the benefits, potentially resulting in comparable or even diminished performance relative to simpler models such as SDPRx. Furthermore, explicitly accounting for environmental covariance was shown to further enhance prediction precision in the presence of overlapping samples. Overall, our findings demonstrate that PleioSDPR is particularly effective when the auxiliary trait has high genetic correlation, greater heritability, and minimal sample overlap with the target trait.

However, we have identified several limitations that will be addressed in future research. First, PleioSDPR currently requires the estimation of a multitude of parameters, which may increase variance and instability in the model. Future work will involve pinpointing and simplifying components of the model that are unnecessarily complex, potentially reducing the number of parameters needed. Additionally, the PleioSDPR computational process is rather time-consuming, even when leveraging parallel computation across 22 chromosomes with the allocation of six threads per chromosome. To enhance efficiency, we are exploring the possibility of rewriting the code using C++ programming. Lastly, PleioSDPR is limited to analyzing only two correlated traits simultaneously. Moving forward, we will extend the model’s capability to include multiple correlated traits, broadening the scope of our genetic architecture analysis.

## References

1. Khera, A. V. et al. Genome-wide polygenic scores for common diseases identify individuals with risk equivalent to monogenic mutations. Nat Genet 50, 1219–1224 (2018).

2. Visscher, P. M. et al. 10 Years of GWAS Discovery: Biology, Function, and Translation. Am J Hum Genet 101, 5–22 (2017).

3. Abdellaoui, A., Yengo, L., Verweij, K. J. H. & Visscher, P. M. 15 years of GWAS discovery: Realizing the promise. The American Journal of Human Genetics 110, 179–194 (2023).

4. Zhang, Y. et al. SUPERGNOVA: local genetic correlation analysis reveals heterogeneous etiologic sharing of complex traits. Genome Biol 22, 262 (2021).

5. Zhang, C., Zhang, Y., Zhang, Y. & Zhao, H. Benchmarking of local genetic correlation estimation methods using summary statistics from genome-wide association studies. Briefings in Bioinformatics 24, bbad407 (2023).

6. Bulik-Sullivan, B. et al. An atlas of genetic correlations across human diseases and traits. Nat Genet 47, 1236–1241 (2015).

7. Shi, H., Mancuso, N., Spendlove, S. & Pasaniuc, B. Local Genetic Correlation Gives Insights into the Shared Genetic Architecture of Complex Traits. The American Journal of Human Genetics 101, 737–751 (2017).

8. Werme, J., van der Sluis, S., Posthuma, D. & de Leeuw, C. A. An integrated framework for local genetic correlation analysis | Nature Genetics. Nature Genetics 54, 274–282 (2022).

9. Stearns, F. W. One Hundred Years of Pleiotropy: A Retrospective. Genetics 186, 767–773 (2010).

10. Ruan, Y. et al. Improving Polygenic Prediction in Ancestrally Diverse Populations. medRxiv (2021) doi:10.1101/2020.12.27.20248738.

11. Ge, T., Chen, C.-Y., Ni, Y., Feng, Y.-C. A. & Smoller, J. W. Polygenic prediction via Bayesian regression and continuous shrinkage priors. Nat Commun 10, 1776 (2019).

12. Zhou, G., Chen, T. & Zhao, H. SDPRX: A statistical method for cross-population prediction of complex traits. The American Journal of Human Genetics 110, 13–22 (2023).

13. Hu, Y. et al. Joint modeling of genetically correlated diseases and functional annotations increases accuracy of polygenic risk prediction. PLOS Genetics 13, e1006836 (2017).

14. Xu, C., Ganesh, S. K. & Zhou, X. mtPGS: Leverage multiple correlated traits for accurate polygenic score construction. The American Journal of Human Genetics (2023) doi:10.1016/j.ajhg.2023.08.016.

15. Gao, B., Yang, C., Liu, J. & Zhou, X. Accurate genetic and environmental covariance estimation with composite likelihood in genome-wide association studies. PLOS Genetics 17, e1009293 (2021).

16. Xiao, J. et al. XPXP: improving polygenic prediction by cross-population and cross-phenotype analysis. Bioinformatics 38, 1947–1955 (2022).

17. Gibbs Sampling Methods for Stick-Breaking Priors: Journal of the American Statistical Association: Vol 96, No 453. https://www.tandfonline.com/doi/abs/10.1198/016214501750332758.

18. Zhai, S., Zhang, H., Mehrotra, D. V. & Shen, J. Pharmacogenomics polygenic risk score for drug response prediction using PRS-PGx methods. Nat Commun 13, 5278 (2022).

19. Berisa, T. & Pickrell, J. K. Approximately independent linkage disequilibrium blocks in human populations. Bioinformatics 32, 283–285 (2016).

20. LDpred2: better, faster, stronger | Bioinformatics | Oxford Academic. https://academic.oup.com/bioinformatics/article/36/22-23/5424/6039173.

21. Bycroft, C. et al. The UK Biobank resource with deep phenotyping and genomic data. Nature 562, 203–209 (2018).

22. Yang, J., Lee, S. H., Goddard, M. E. & Visscher, P. M. GCTA: A Tool for Genome-wide Complex Trait Analysis. Am J Hum Genet 88, 76–82 (2011).

23. Second-generation PLINK: rising to the challenge of larger and richer datasets | GigaScience | Oxford Academic. https://academic.oup.com/gigascience/article/4/1/s13742-015-0047-8/2707533?login=false.

24. Purcell, S. & Chang, C. PLINK 2.0.

25. Loh, P.-R. et al. Efficient Bayesian mixed-model analysis increases association power in large cohorts. Nat Genet 47, 284–290 (2015).

26. Bulik-Sullivan, B. K. et al. LD Score regression distinguishes confounding from polygenicity in genome-wide association studies. Nat Genet 47, 291–295 (2015).

27. New genetic loci link adipose and insulin biology to body fat distribution | Nature. https://www.nature.com/articles/nature14132.

28. Gaulton, K. J. et al. Genetic fine mapping and genomic annotation defines causal mechanisms at type 2 diabetes susceptibility loci. Nat Genet 47, 1415–1425 (2015).

29. Clarke, L. et al. The international Genome sample resource (IGSR): A worldwide collection of genome variation incorporating the 1000 Genomes Project data. Nucleic Acids Res 45, D854–D859 (2017).

30. Xu, L., Zhou, G., Jiang, W. et al. JointPRS: A data-adaptive framework for multi-population genetic risk prediction incorporating genetic correlation. Nat Commun 16, 3841 (2025). 10.1038/s41467-025-59243-x

