## Supplementary figures and tables for "Joint Modeling of Effect Sizes for Two Correlated Traits: Characterizing Trait Properties to Enhance Polygenic Risk Prediction"

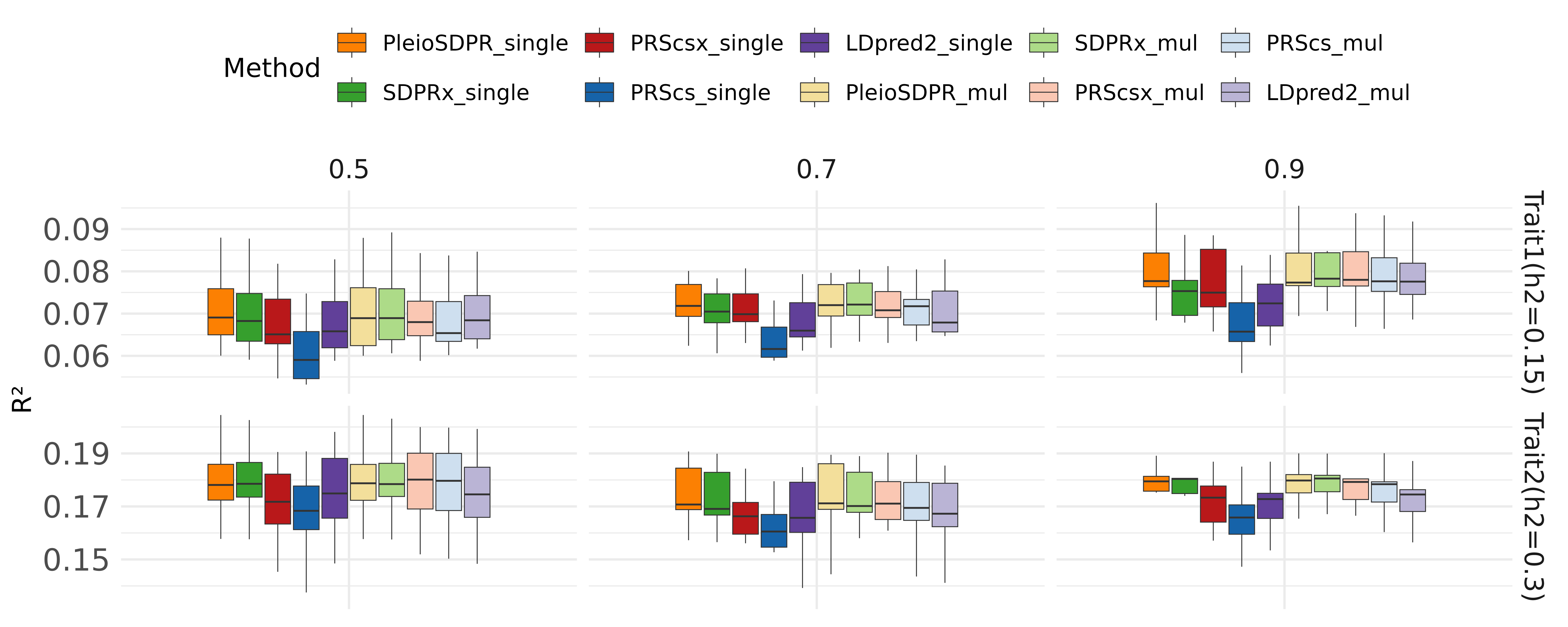

**Supplementary Figure 1: Simulation results when there is partial sample overlap.** We compare the methods considering with validation dataset and without validation dataset. When there is no validation dataset, we name them as PleioSDPR_single, SDPRx_single, PRScsx_single, PRScs_single and LDpred2_single. When there is a validation dataset, we name them PleioSDPR_mul, SDPRx_mul, PRScsx_mul, PRScs_mul, and LDpred2_mul.

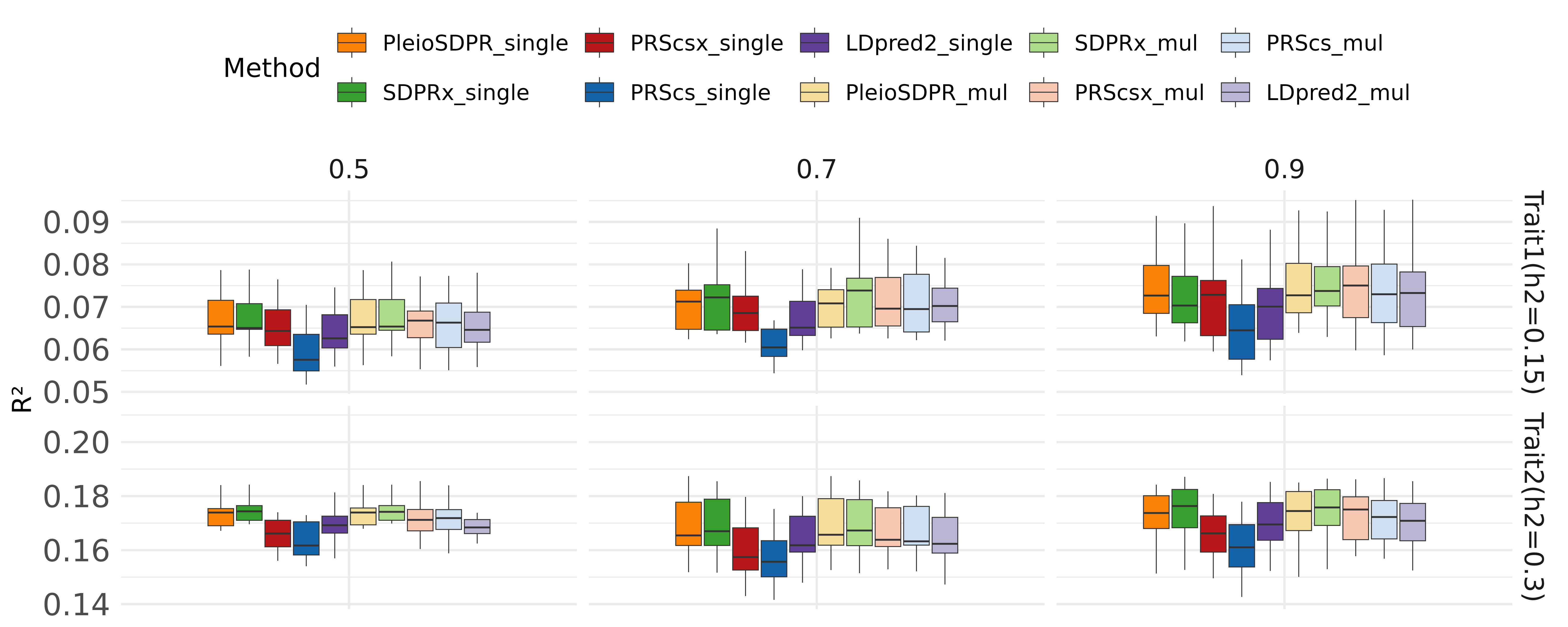

**Supplementary Figure 2: Simulation results when there is complete sample overlap.** We compare the methods considering with validation dataset and without validation dataset. When there is no validation dataset, we name them as PleioSDPR_single, SDPRx_single, PRScsx_single, PRScs_single and LDpred2_single. When there is a validation dataset, we name them PleioSDPR_mul, SDPRx_mul, PRScsx_mul, PRScs_mul, and LDpred2_mul.

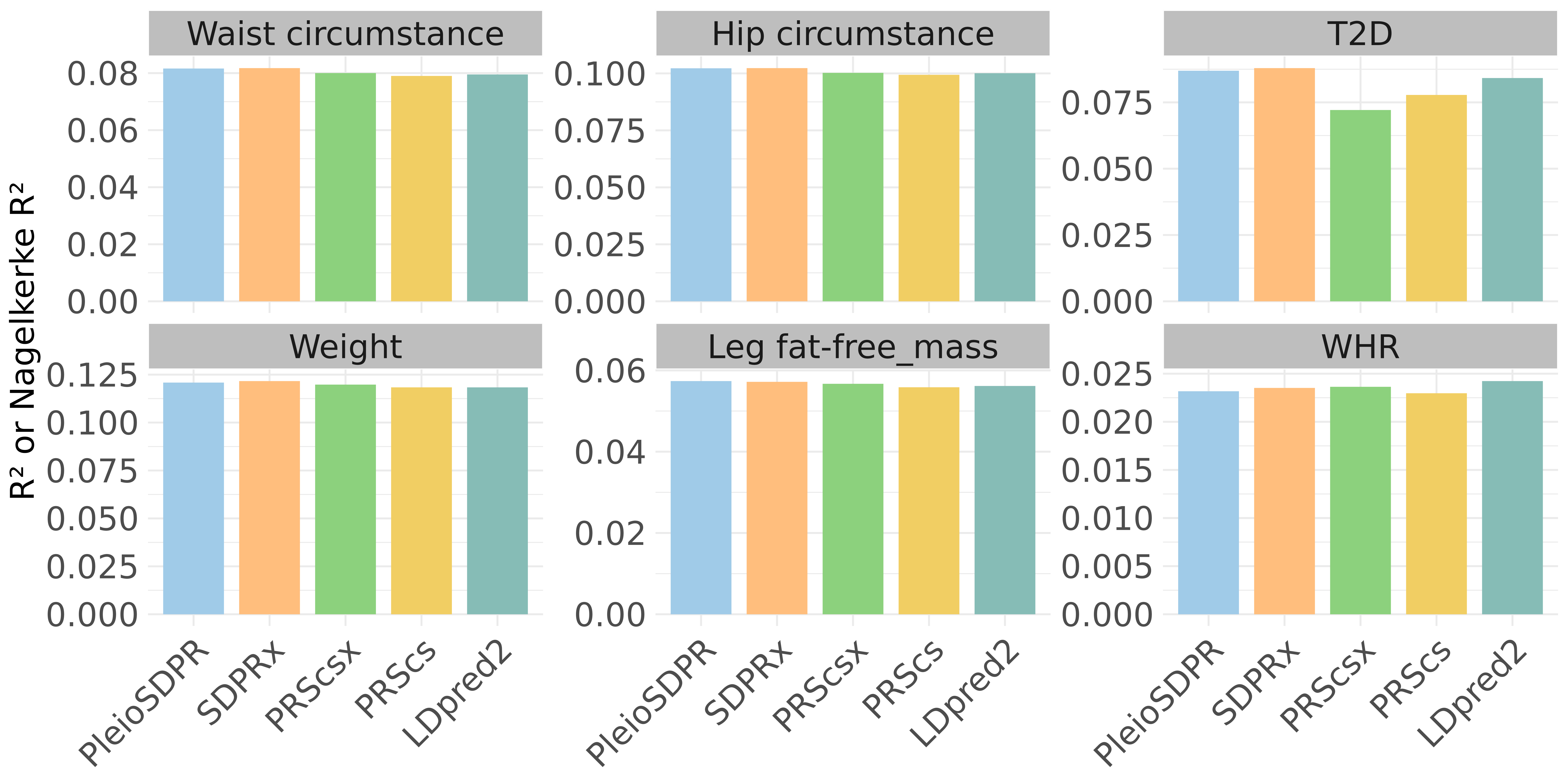

**Supplementary Figure 3: Prediction performance of different methods for 3 trait pairs that have no sample overlap when there is a validation dataset.**

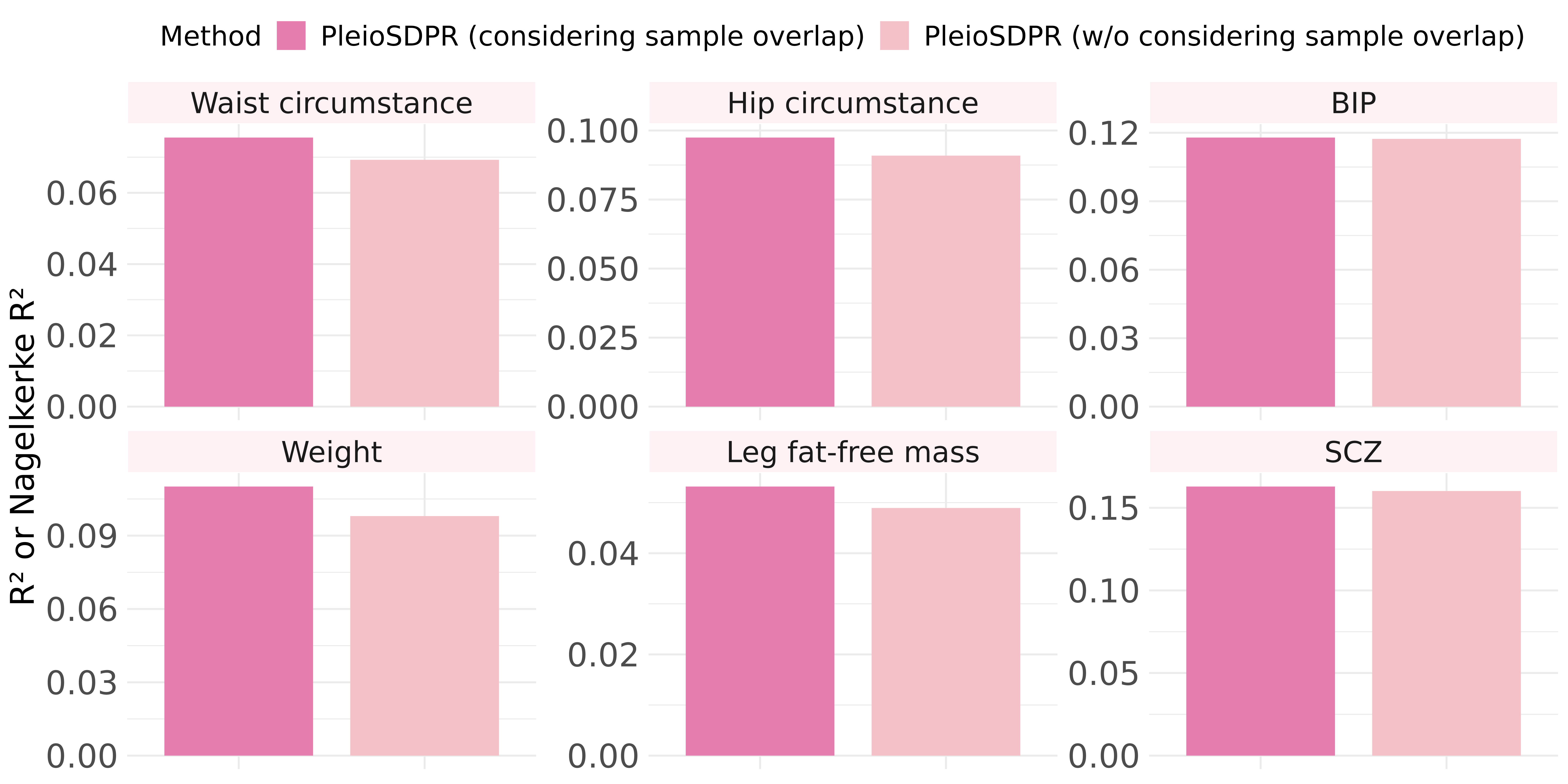

**Supplementary Figure 4: The comparison of PleioSDPR before and after considering sample overlap in its model when two traits have partial sample overlap.**

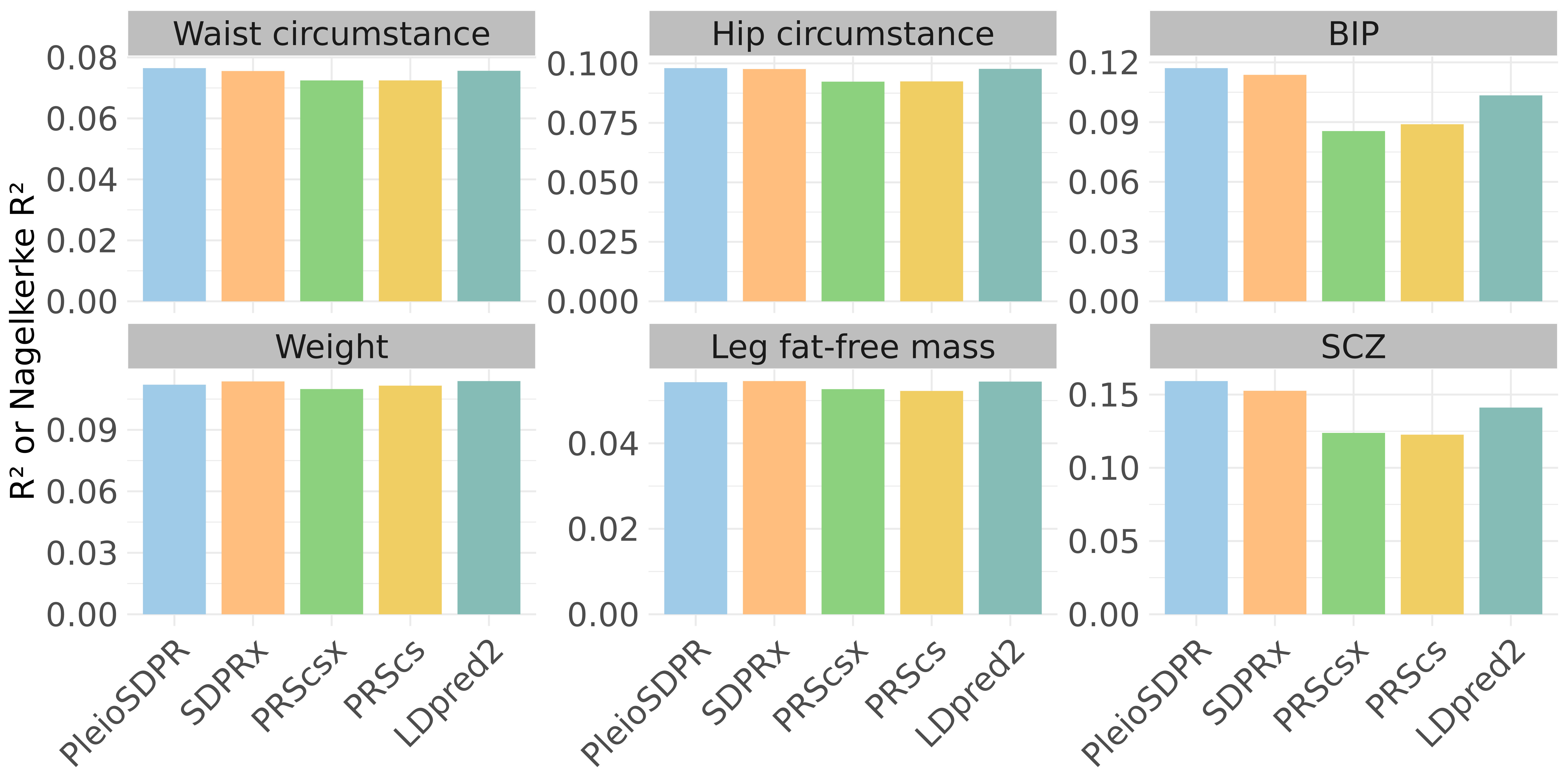

**Supplementary Figure 5: Prediction performance of different methods for 3 trait pairs that have partial sample overlap when there is a validation dataset.**

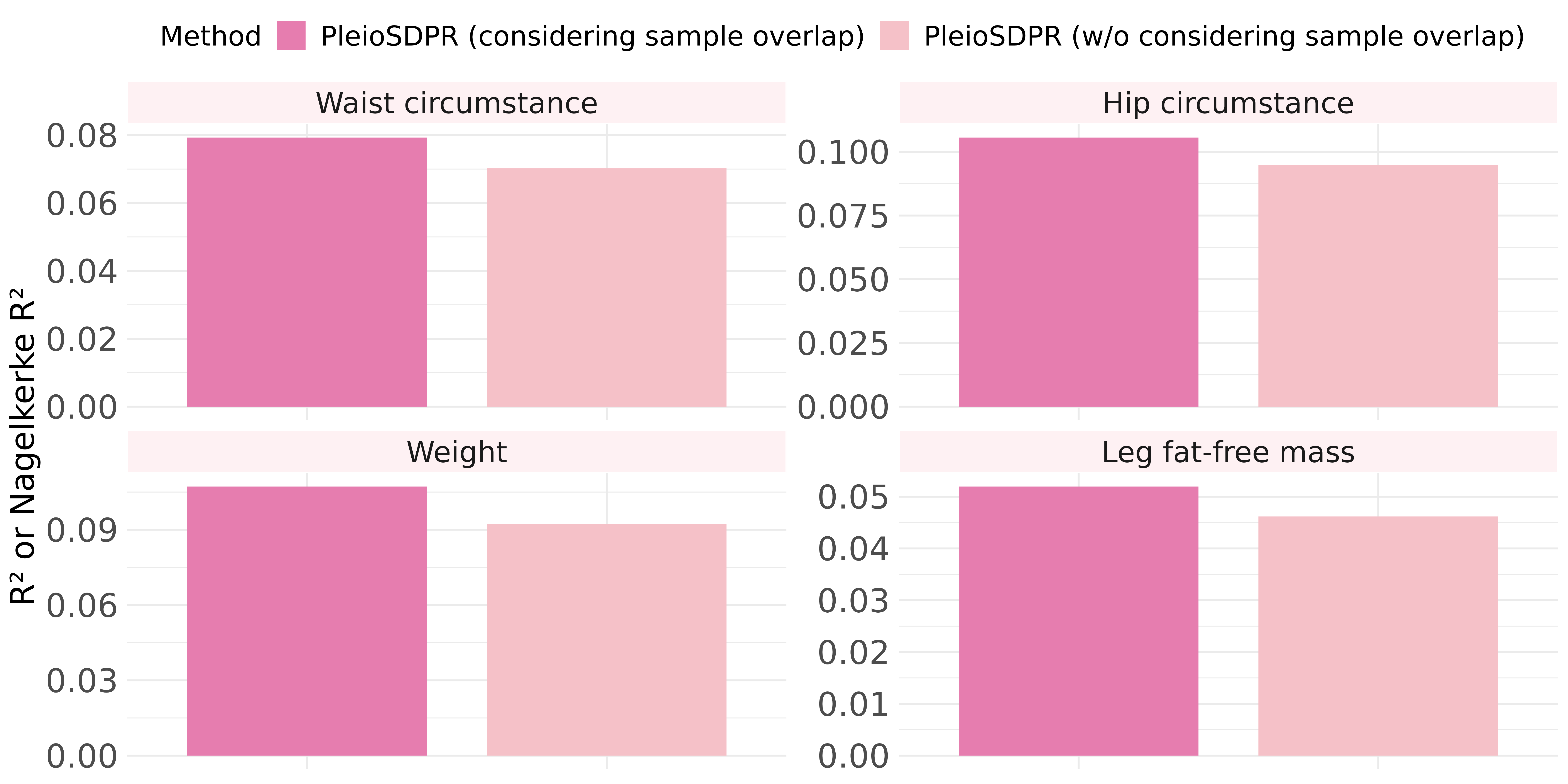

**Supplementary Figure 6: The comparison of PleioSDPR before and after considering sample overlap in its model when two traits have partial sample overlap.**

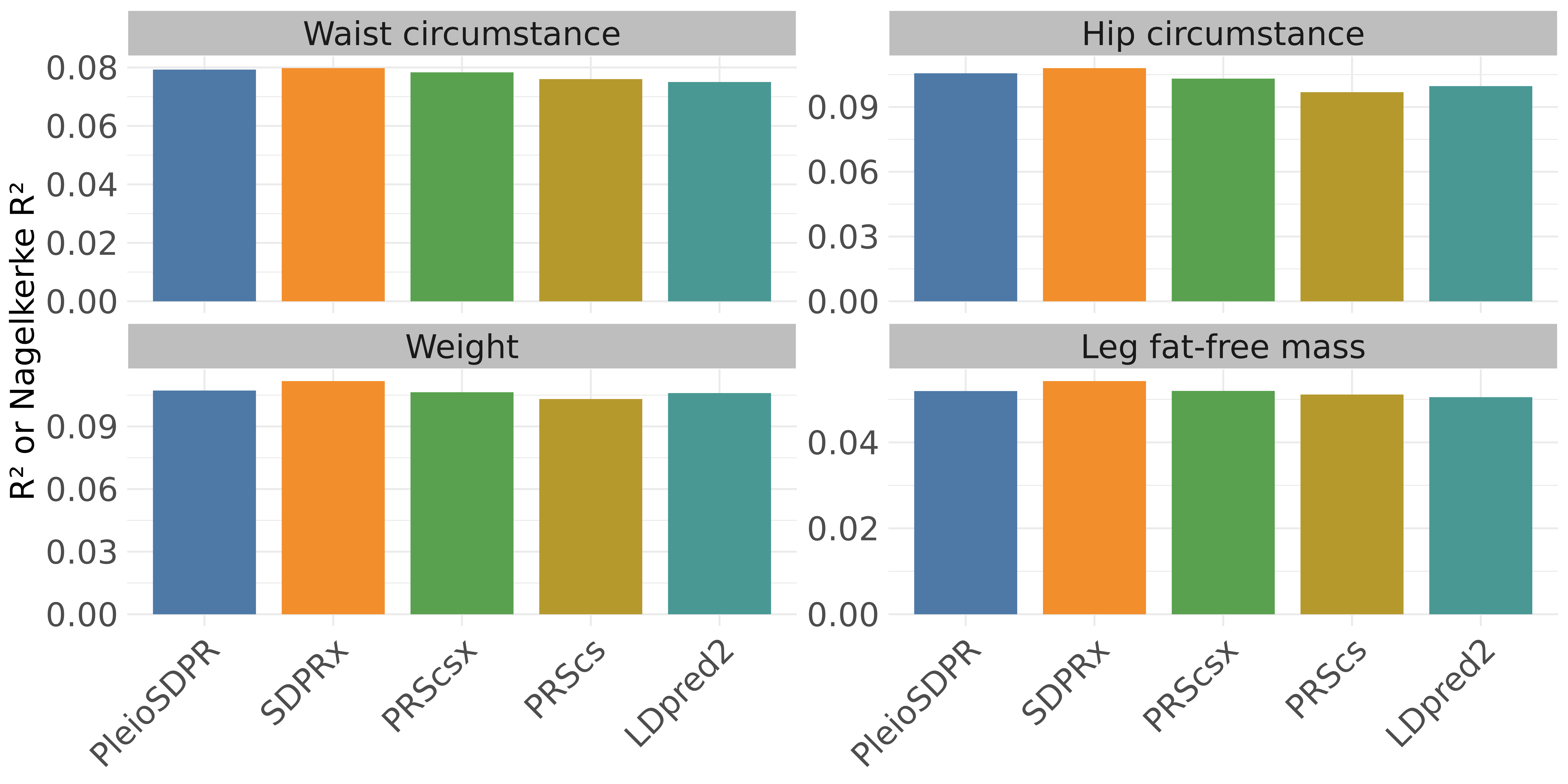

**Supplementary Figure 7: Prediction performance of different methods for 2 trait pairs that have complete sample overlap when there is no validation dataset.**

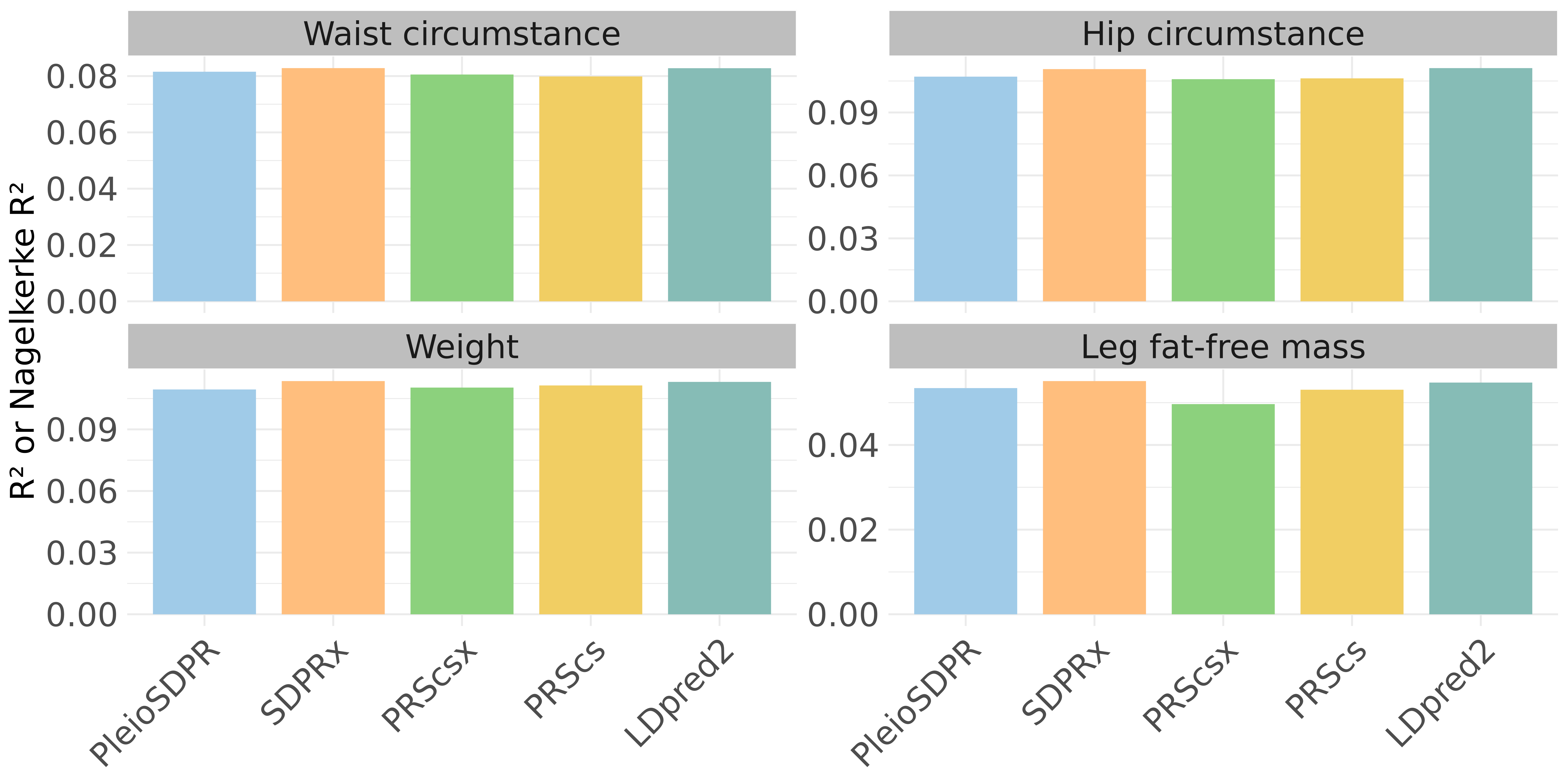

**Supplementary Figure 8Prediction performance of different methods for 2 trait pairs that have complete sample overlap when there is a validation dataset.**

### **Supplementary Tables**

| **_Trait_** | **_Validation dataset 1_** | **_Validation dataset 2_** | **_Validation dataset 3_** | **_Testing dataset_** |
| --- | --- | --- | --- | --- |
| _Waist circumstance_ | _1,638_ | _5,236_ | _8,058_ | _61,039_ |
| _Hip circumstance_ | _1,638_ | _5,236_ | _8,053_ | _61,034_ |
| _T2D_ | _1,261_ | _4,155_ | _6,341_ | _51,489_ |
| _Weight_ | _1,637_ | _5,233_ | _8,038_ | _60,975_ |
| _Leg fat-free mass_ | _1,618_ | _5,155_ | _7,931_ | _60,073_ |
| _WHR_ | _1,638_ | _5,235_ | _8,053_ | _61,027_ |

**Supplementary Table 1: Sample sizes of testing and validation datasets when two traits in a trait pair have no sample overlaps.**

| **_Trait_** | **_Validation dataset 1_** | **_Validation dataset 2_** | **_Validation dataset 3_** | **_Testing dataset_** |
| --- | --- | --- | --- | --- |
| _Waist circumstance_ | _1,638_ | _5,236_ | _8,058_ | _61,039_ |
| _Hip circumstance_ | _1,638_ | _5,236_ | _8,053_ | _61,034_ |
| _BIP_ | _/_ | _205_ | _242_ | _2,245_ |
| _Weight_ | _1,637_ | _5,233_ | _8,038_ | _60,975_ |
| _Leg fat-free mass_ | _1,618_ | _5,155_ | _7,931_ | _60,073_ |
| _SCZ_ | _/_ | _123_ | _246_ | _1,060_ |

**Supplementary Table 2: Sample sizes of testing and validation datasets when two traits in a trait pair have partial samples overlapped.**
