## Supplementary method for "Joint Modeling of Effect Sizes for Two Correlated Traits: Characterizing Trait Properties to Enhance Polygenic Risk Prediction"

### **Supplementary Methods**

**S1. Likelihood Function**

When there are two GWASs, with $N_{1}$ samples for trait 1 ($Y_{1})$, $N_{2}$ samples for trait 2 ($Y_{2}$) and $N_{s}$ overlapped samples, we assume that:

|  | $Y_{1}=X_{1}\beta_{1}+\epsilon_{1}$ $Y_{2}=X_{2}\beta_{2}+\epsilon_{2}$ | ((1) |
| --- | --- | --- |

where $X_{1}$ and $X_{2}$ are the standardized genotypes for $N_{1}$and $N_{2}$samples, $\beta_{1}$ and $\beta_{2}$ are the true effect size vectors for two traits, and $\epsilon_{1}$ and $\epsilon_{2}$ are the vectors of noises with $var\left( \epsilon_{1} \right)=1-h_{1}^{2}$ and $var\left( \epsilon_{2} \right)=1-h_{2}^{2}$, $h_{1}^{2}$ and $h_{2}^{2}$ are heritability for two traits, respectively. We futher assume that the first $N_{s}$samples overlap with

|  | $cov\left( \epsilon_{1,j_{1}}, \epsilon_{2,j_{2}} \right)=\left\{ \begin{aligned} \rho_{e},1\leq j_{1}=j_{2}\leq N_{s} \\ 0, otherwise \end{aligned} \right.,$ | ((2) |
| --- | --- | --- |

where $\rho_{e}$ is the environmental covariance.

Since there are possible samples overlapped between two complex traits in one population, the conditional covariance of marginal effect sizes is no longer zero since

$$Cov\left( \hat{\beta}_{1},\hat{\beta}_{2} | \beta_{1},\beta_{2} \right)=E\left( \hat{\beta}_{1}\hat{\beta}_{2}^{T} | \beta_{1},\beta_{2} \right)-E\left( \hat{\beta}_{1} | \beta_{1} \right)E\left( \hat{\beta}_{2} | \beta_{2} \right)^{T}$$

$$=\frac{1}{N_{1}N_{2}}E\left( X_{1}\left( X_{1}^{T}\beta_{1}+\epsilon_{1} \right){(X}_{2}\left( X_{2}^{T}\beta_{2}+\epsilon_{2} \right))T |\beta_{1},\beta_{2} \right)-\left( D\beta_{1} \right)\left( D\beta_{2} \right)^{T}$$

$$=\frac{1}{N_{1}N_{2}}X_{1}E\left( \epsilon_{1}\epsilon_{2}^{T} | \beta_{1},\beta_{2} \right)X_{2}^{T}$$

$$=\frac{\rho_{e}}{N_{1}N_{2}}X^{i*}X^{i*T}$$

$$=\frac{N_{s}\rho_{e}}{N_{1}N_{2}D} .$$

(3)

Thus, the likelihood connects the pair of marginal effect sizes in GWAS summary statistics for two traits with true effect sizes through a multivariate normal distribution accounting for LD is changed to be

$${\hat{\beta}_{1}| \atop\hat{\beta}_{2}|}{\beta_{1} \atop\beta_{2}}\sim N\left( \binom{D\beta_{1}}{D\beta_{2}},\left( \begin{matrix} D/N_{1} & N_{s}\rho_{e}D/N_{1}N_{2} \\ N_{s}\rho_{e}D/N_{1}N_{2} & D/N_{2} \end{matrix} \right) \right).$$

(4)

**S2 Modified Likelihood Function**

Besides, since individual-level data is difficult to obtain, external reference panels are typically used to estimate LD matrices, like data from 1000 Genomes Project and UK Biobank. The effect sizes of SNPs in summary statistic may diverge from the expected values based on the likelihood function (4) and the external reference LD matrix, especially for SNPs genotyped on different sets of samples and in strong LD. If SNPs for GWAS of one trait are genotyped on different individuals, then we can modify the likelihood function (4) into:

$${\hat{\beta}_{1}| \atop\hat{\beta}_{2}|}{\beta_{1} \atop\beta_{2}}\sim N\left( \binom{D\beta_{1}}{D\beta_{2}},\left( \begin{matrix} (D+aI)/N_{1} & N_{s}\rho_{e}(D+aI)/N_{1}N_{2} \\ N_{s}\rho_{e}(D+aI)/N_{1}N_{2} & (D+aI)/N_{2} \end{matrix} \right) \right).$$

(5)

where a is set to be 0.1.

**S3 Prior**

For each SNP j, we specify the following joint distribution as the prior on the effect sizes $\beta_{j1}$ and $\beta_{j2}$ for two traits.

|  | 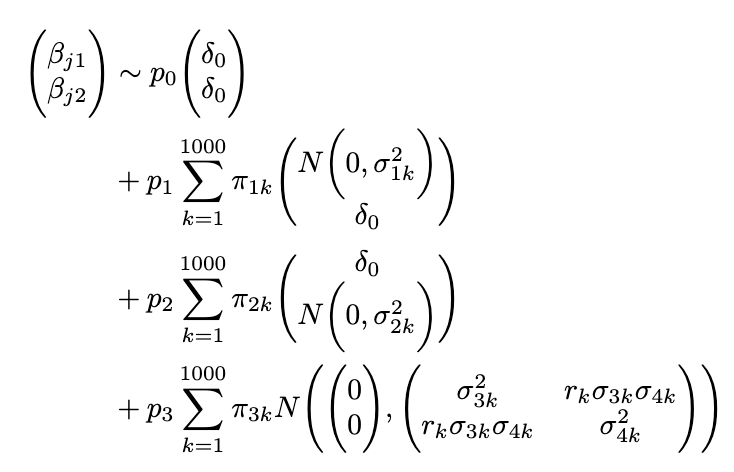 |
| --- | --- |

PleioSDPR adopts the idea of Bayesian nonparametric prior from SDPRx, which used the truncated stick-breaking process to represent the probability of assignments for the second (trait 1 specific), third (trait 2 specific), and fourth terms (shared with correlation). Same as SDPRx, we set the maximum components to 1,000. Thus, we have

|  | 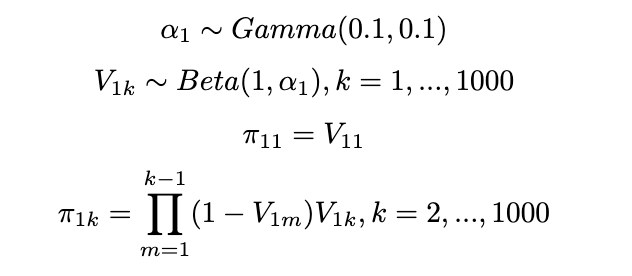, | (6) |
| --- | --- | --- |

|  | 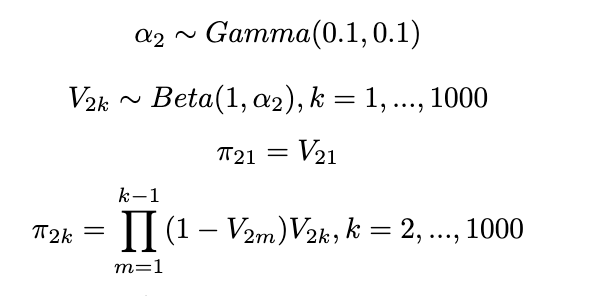, | (7) |
| --- | --- | --- |
|  | 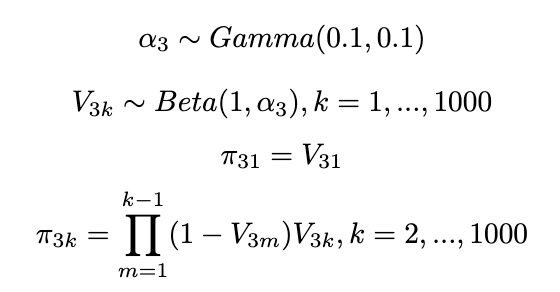. | (8) |

Different from SDPRx, we relax the assumption that the variances are the same in the fourth term and allow the correlations different. Thus, we utilize a hierarchical half-t prior for the variance-covariance matrix:

|  | 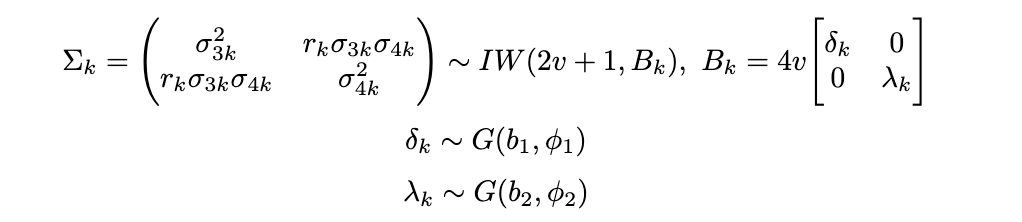. | (9) |
| --- | --- | --- |

For the variances in the second and third terms, we had

|  | 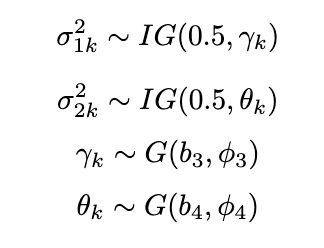. | (10) |
| --- | --- | --- |

We set a Dirichlet distribution prior on the probability of each SNP,

|  | 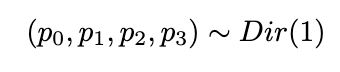. | (11) |
| --- | --- | --- |

**S4 MCMC algorithm**

We partition the matrix using the block matrix multiplication rules to derive the inverse of

|  | 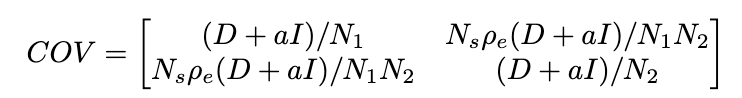. |
| --- | --- |

We set $s=\frac{N_{s}\rho_{e}}{\sqrt{N_{1}N_{2}}}$ as a parameter capturing information of overlapped sample size and environmental covariance, the covariance matrix of likelihood function can be expressed as:

|  | 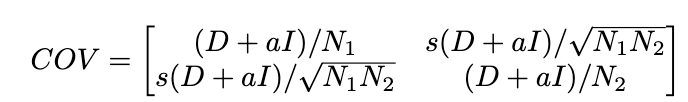. |
| --- | --- |

Then the inverse of the covariance matrix of likelihood function can be derived as:

|  | 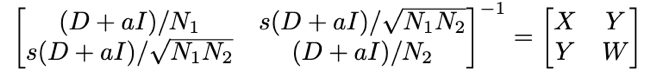. |
| --- | --- |

where $X=\frac{N_{1}}{1-s^{2}}\left( D+aI \right)^{-1}$, $Y=-\frac{s\sqrt{N_{1}N_{2}}}{1-s^{2}}\left( D+aI \right)^{-1}$ and $W=\frac{N_{2}}{1-s^{2}}\left( D+aI \right)^{-1}$.

Then we assign

|  | 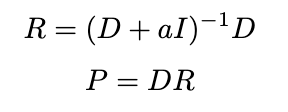. | (12) |
| --- | --- | --- |

and

|  | 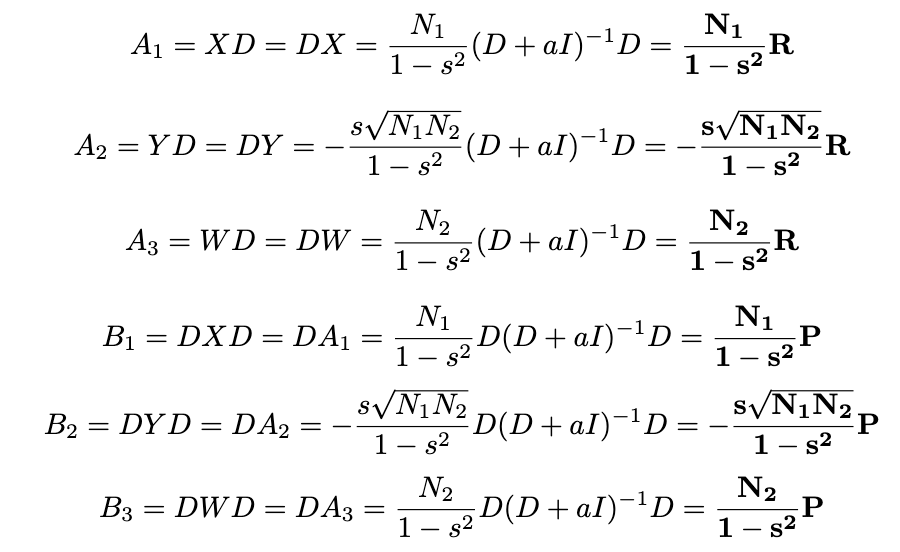. | (13) |
| --- | --- | --- |

**S4.1 Sampling the assignment of SNPs,** $\boldsymbol{z}_{\boldsymbol{j}}$

$z_{j}=(1,k)$ means the SNP j is only the causal SNP for trait 1 and is assigned to the kth component of that category. $z_{j}=(2,k)$ means the SNP j is only the causal SNP for trait 2 and is assigned to the kth component of that category. $z_{j}=(3,k)$ means the SNP j is causal for both traits and is assigned to the kth component of that category. $z_{j}=(0,0)$ means the effect sizes of SNP j are zero for both traits.

We first set

|  | 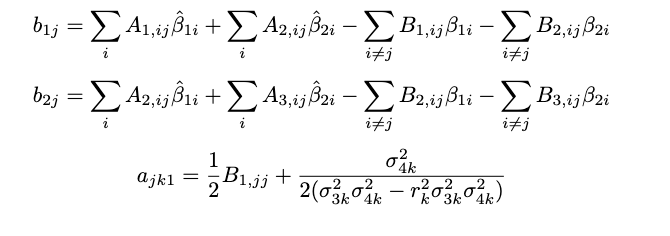  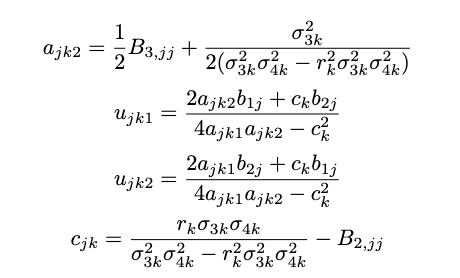 |
| --- | --- |

Then we can derive the posterior probability of each SNP’s assignment in (3,k) as:

|  | .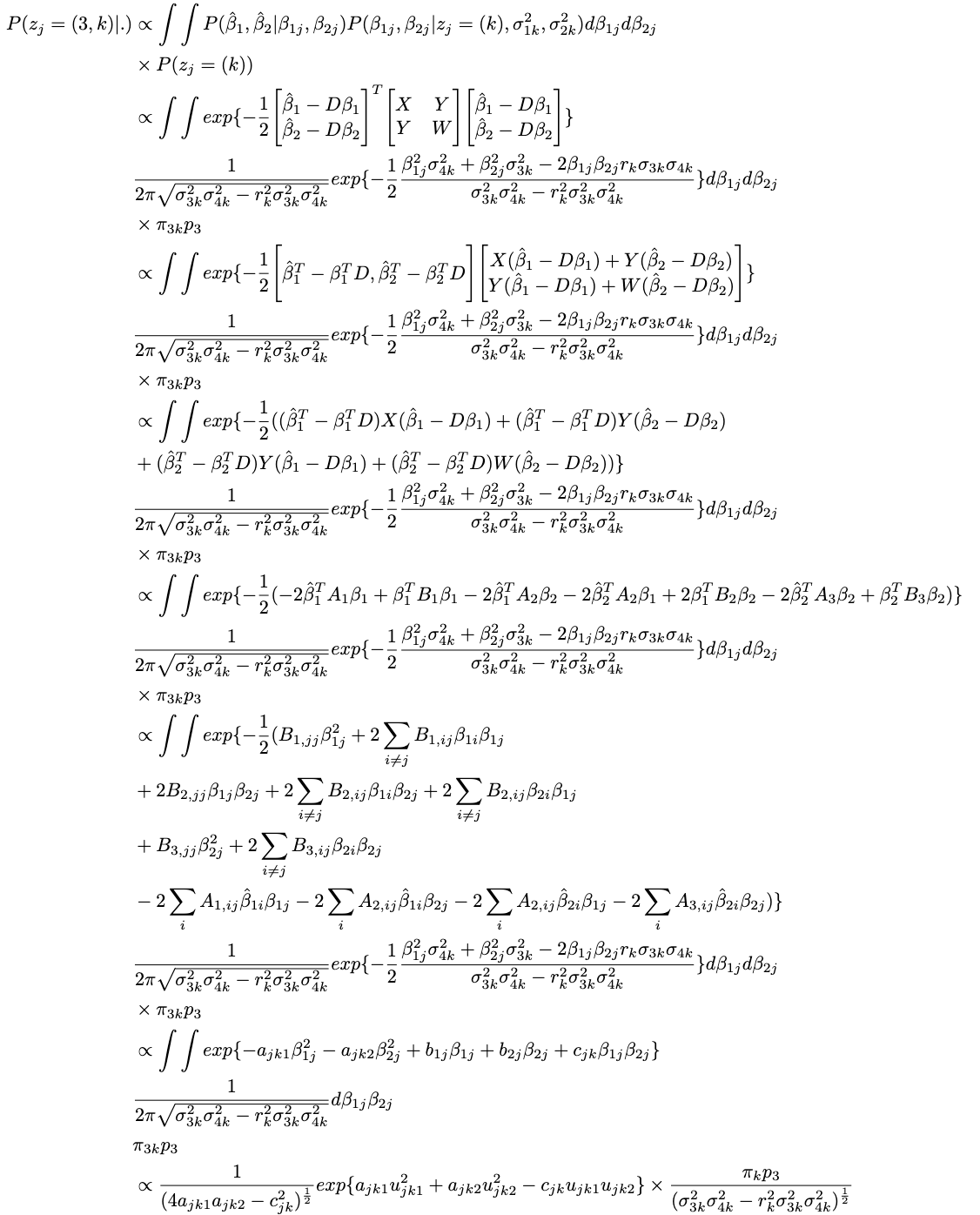 |
| --- | --- |

We next derive the conditional probability of SNP j whose effect sizes are population-specific or null, which are the same as SDPRx.

|  | 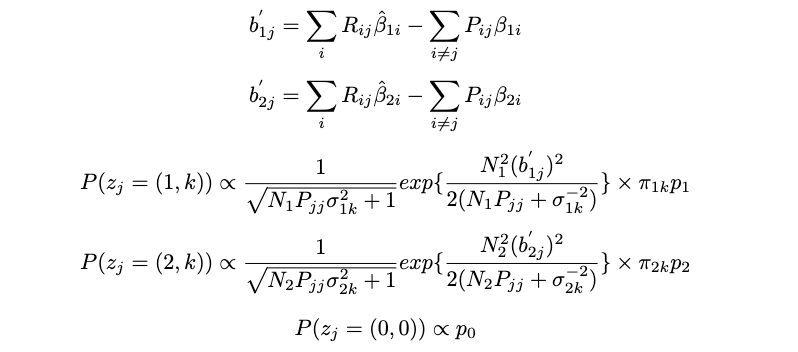 |
| --- | --- |

Based on SDPRx, we use the log-exp-sum trick to avoid numerical overflow. Note that because SNPs in different LD blocks are approximately independent, we can sample their assignments in parallel. For trait 1 specific SNPs, we only need to evaluate $P(z_{j}=(1,k))$ and $P(z_{j}=(0,0))$, and for trait 2 specific SNPs, we only need to evaluate $P(z_{j}=(2,k))$ and $P(z_{j}=(0,0))$.

**S4.2 Sampling the true effect sizes,** $\boldsymbol{\beta}_{\boldsymbol{1}}$ **and** $\boldsymbol{\beta}_{\boldsymbol{2}}$

We jointly sample the effect sizes of causal SNPs in one independent LD block. First, we introduce two indexes $\gamma_{1}$ and $\gamma_{2}$such that $\beta_{1,\gamma_{1}}$ and $\beta_{2,\gamma_{2}}$ are non-zero. We combine $\beta_{1,\gamma_{1}}$ and $\beta_{2,\gamma_{2}}$ into one vector $\beta_{\gamma}$ which follows a bivariate normal distribution with mean 0 and variance-covariance matrix $\Sigma_{0}.$ For example that we assume that $\gamma_{1}=\gamma_{2}=\left\{ 1,2,3 \right\}$, so we can obtain that $\beta_{\gamma}=(\beta_{1,1}, \beta_{1,2}, \beta_{1,3}, \beta_{2,1}, \beta_{2,2}, \beta_{2,3})$, thus

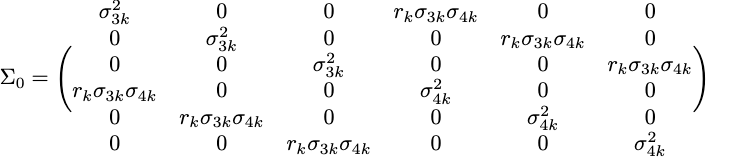
.

We can further derive the posterior probability of the true effect sizes as:

|  | .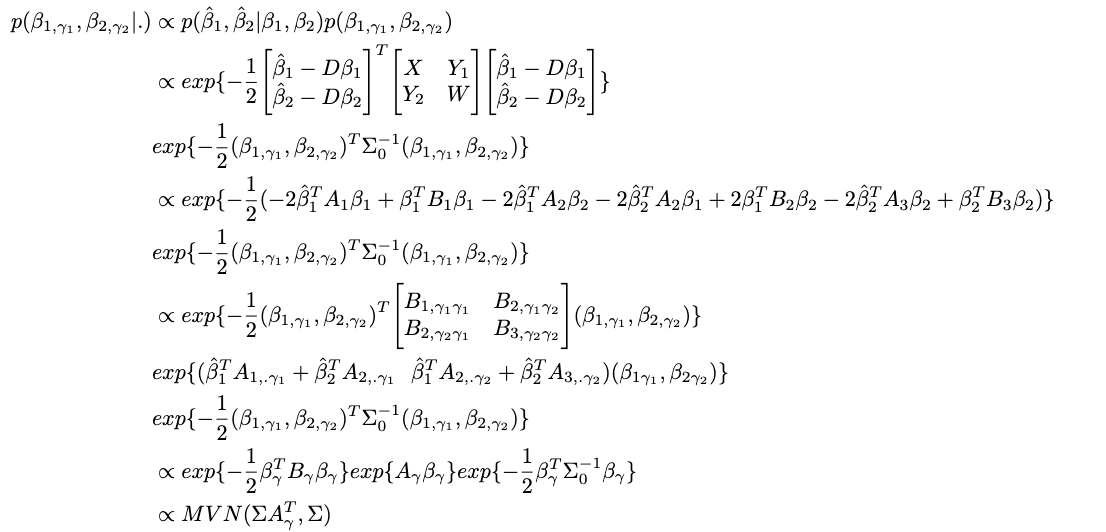 | (14) |
| --- | --- | --- |

where

|  | 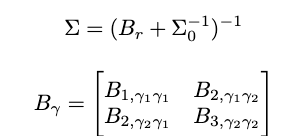 |
| --- | --- |

with dimension $\left( \gamma_{1}+\gamma_{2} \right)$ by $\left( \gamma_{1}+\gamma_{2} \right)$.

|  | 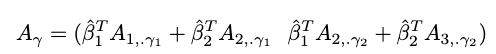 |
| --- | --- |

with dimension 1 by $\left( \gamma_{1}+\gamma_{2} \right)$.

Besides, $A_{1,\gamma_{2}}$ and $A_{2,\gamma_{2}}$ are two sub-matrices of $A_{1}$ and $A_{2}$ where only selecting columns with indexes $\gamma_{1}$ and $\gamma_{2}$. $B_{1,\gamma_{1}\gamma_{1}}$, $B_{2,\gamma_{1}\gamma_{2}}$, $B_{2,\gamma_{2}\gamma_{1}}$ and $B_{3,\gamma_{2}\gamma_{2}}$are four sub-matrices of $B_{1}$, $B_{2}$ and $B_{3}$ where selecting both rows and columns with relative indexes $\gamma_{1}$ or $\gamma_{2}$.

**S4.3 Sampling variances for trait-specific effects,** $\boldsymbol{\sigma}_{\boldsymbol{1}\boldsymbol{k}}^{\boldsymbol{2}}$ **and** $\boldsymbol{\sigma}_{\boldsymbol{2}\boldsymbol{k}}^{\boldsymbol{2}}$

We assume that $\sigma_{1k}^{2}$follows $IG(0.5,\gamma_{k})$ and $\sigma_{2k}^{2}$ follows $IG(0.5,\theta_{k})$ where $\gamma_{k}\sim G(b_{3},\phi_{3})$ and $\theta_{k}\sim G\left( b_{4}, \phi_{4} \right).$ Thus we can obtain the posterior probability of $\sigma_{1k}^{2}$ and $\sigma_{2k}^{2}:$

|  | 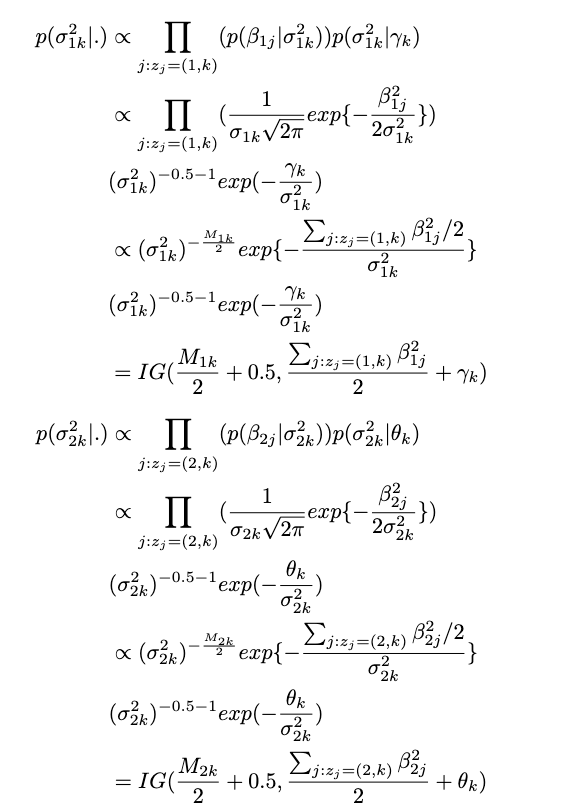 | (15) |
| --- | --- | --- |

*Then we obtain the posterior distribution of* $\gamma_{k}$ *and* $\theta_{k}$*.*

|  | 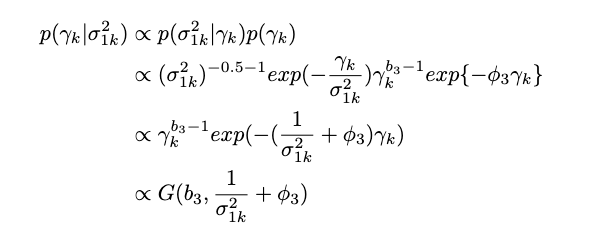  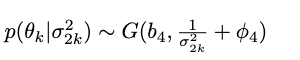 | (16) |
| --- | --- | --- |

**S4.4 Sampling the variance-covariance matrix for effects shared between traits,** $\boldsymbol{\sigma}_{\boldsymbol{3}\boldsymbol{k}}^{\boldsymbol{2}}$

When $z_{j}=(3,k)$, $\binom{\beta_{1j}}{\beta_{2j}}\sim N(\binom{0}{0},\Sigma_{k})$, where $\Sigma_{k}=\left( {\sigma_{3k}^{2} \atop r_{k}\sigma_{3k}\sigma_{4k}}{r_{k}\sigma_{3k}\sigma_{4k} \atop\sigma_{4k}^{2}} \right)$. We assume $\Sigma_{k}\sim IW\left( 2w+1, B_{k} \right), B_{k}=4v\left[ \begin{matrix} \delta_{k} & 0 \\ 0 & \lambda_{k} \end{matrix} \right], \delta_{k}\sim G\left( b_{1},\phi_{1} \right),\lambda_{k}\sim G\left( b_{2},\phi_{2} \right).$ Thus the posterior distribution of $\Sigma_{k}$ is:

|  | 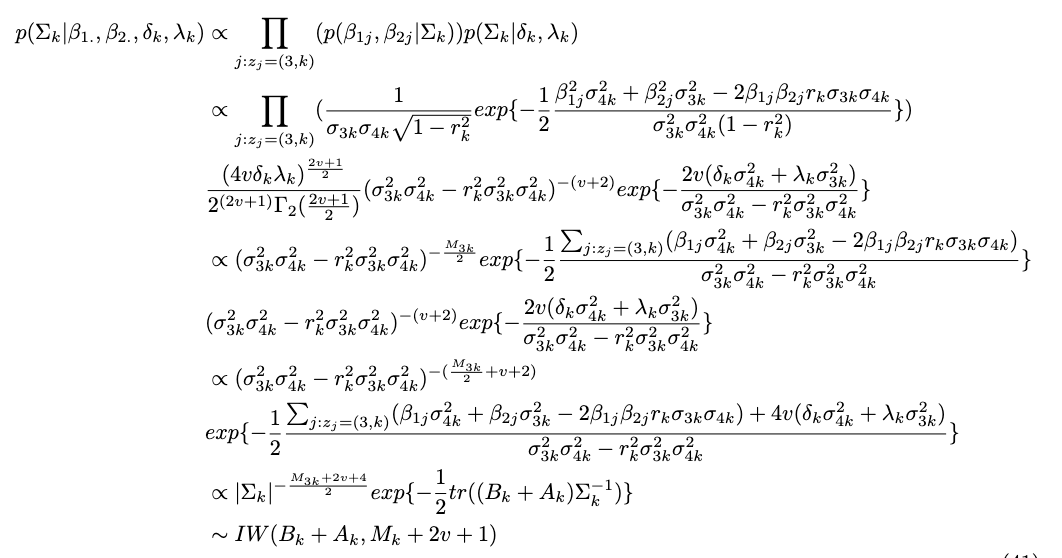  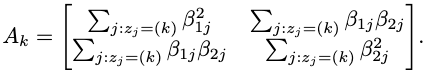 | (17) |
| --- | --- | --- |

Then we can derive the posterior distribution of $\delta_{k}$ and $\lambda_{k}$ as

|  | 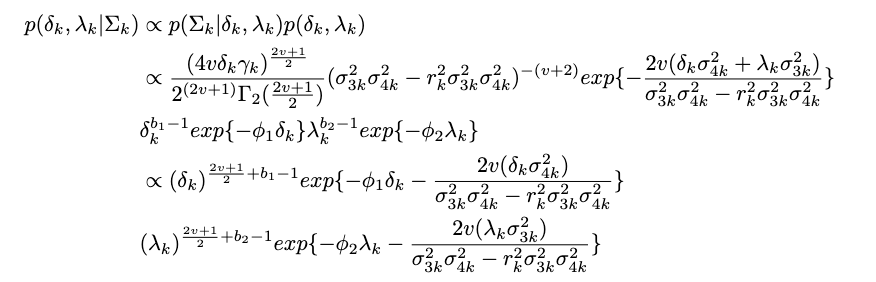. |
| --- | --- |

Thus

|  | 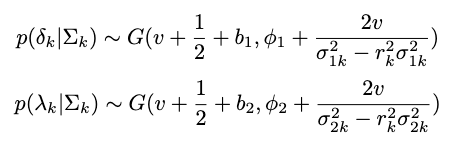. | (18) |
| --- | --- | --- |

And we set $b_{1}=b_{2}=0.5$, $\phi_{1}=\phi_{2}=1$ and $v=0.5$.

**S4.5 Sampling** $\boldsymbol{v}_{\boldsymbol{mk}}$

$m\in\left\{ 1,2,3 \right\}$ *and* $k\in\{1,2,\ldots,1000\}$*. The full conditional likelihood is*

|  | 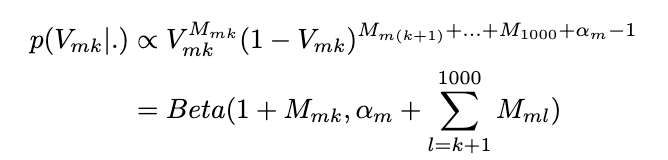. | (19) |
| --- | --- | --- |

for $j=1,2,3$ and $k=1,\ldots,999$. $V_{m,1000}$equals 1 according to the definition of the truncated stick-breaking process.

**S4.6 Sampling** $\boldsymbol{\pi}_{\boldsymbol{mk}}$

The posterior probability of $\pi_{mk}, m\in\{1,2,3\}$ can be computed as

|  | 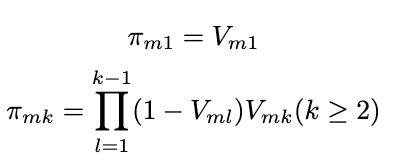. | (20) |
| --- | --- | --- |

**S4.7 Sampling** $\boldsymbol{p}_{\boldsymbol{1}}\boldsymbol{,}\boldsymbol{p}_{\boldsymbol{2}}\boldsymbol{,}\boldsymbol{p}_{\boldsymbol{3}}\boldsymbol{,}\boldsymbol{p}_{\boldsymbol{4}}$

The conditional distribution is

|  | .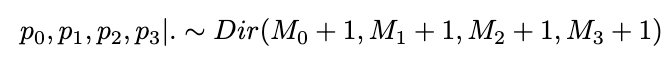 | (21) |
| --- | --- | --- |

where$M_{0} =\sum_{j} I(z_{j}=(0,0)),M_{1} =\sum_{j} I(z_{j}=(1,.)),M_{2} =\sum_{j} I\left( z_{j}=\left( 2,. \right) \right) M_{3} =\sum_{j} I(z_{j}=(3,.)).$Note that we exclude GWAS-specific variants when computing $M_{0}$, $M_{1}$, $M_{2}$, $M_{3}.$

**S4.8 Sampling** $\boldsymbol{\alpha}_{\boldsymbol{m}}$

The full conditional likelihood of $\alpha_{m}$ is

|  | 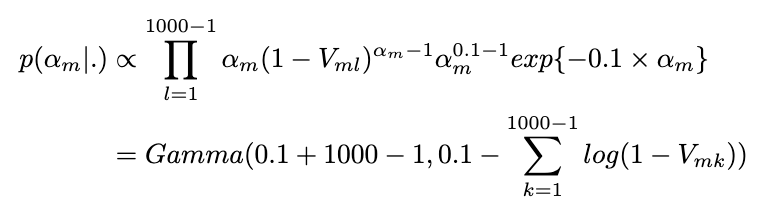. | (22) |
| --- | --- | --- |

**S5 How to obtain** $\boldsymbol{s=}\frac{\boldsymbol{N}_{\boldsymbol{s}}\boldsymbol{\rho}_{\boldsymbol{e}}}{\sqrt{\boldsymbol{N}_{\boldsymbol{1}}\boldsymbol{N}_{\boldsymbol{2}}}}$

We can estimate $\frac{N_{s}\rho_{e}}{\sqrt{N_{1}N_{2}}}$ where $\rho_{t}=\rho_{g}+\rho+e$is the sum of genetic covariance and environmental covariance.

|  | 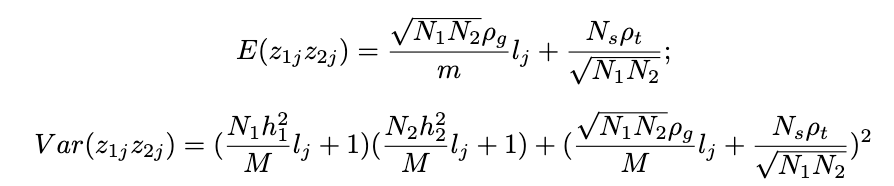. | (23) |
| --- | --- | --- |

where $z_{1j}$ and $z_{2j}$ are the z-scores of the SNP j for triat1 and triat2, respectively, and $l_{j}$ is the LD-score of SNP j. Thus, by regressing the product of of $z_{1j}$ and $z_{2j}$ to the LD score $l_{j}$ using the weighted linear regression, we can obtain the estimated parameter $\hat{\rho}_{g}$ and $\hat{N_{s}\rho_{t}}$from the fitted slope and intercept.
